# Neuroinflammatory Stress Preferentially Impacts Synaptic MAPK Signaling and Mitochondria in Excitatory Neurons

**DOI:** 10.1101/2025.10.03.680087

**Authors:** Claudia Espinosa-Garcia, Upasna Srivastava, Prateek Kumar, Dilpreet Kour, Sneha Malepati, Brendan R. Tobin, Hailian Xiao, Sydney Sunna, Christine A. Bowen, Lihong Cheng, Pritha Bagchi, Duc M. Duong, Ted J. Whitworth, Liu Xinran, Nicholas T. Seyfried, Levi B. Wood, Victor Faundez, Srikant Rangaraju

## Abstract

**Background:** Understanding synapse-specific effects of neuroinflammation can provide mechanistic and therapeutically relevant insights across the spectrum of neurological diseases.

**Methods:** We applied neuron-specific proteomic biotinylation *in vivo*, differential centrifugation of brain for crude synaptosome enrichment (P2 fraction) and mass spectrometry (MS) analysis of biotinylated proteins to derive native-state proteomes of Camk2a-positive neurons and their corresponding P2 synaptic compartments. Next, in an *in vivo* model of systemic lipopolysaccharide (LPS) dosing, we examined the effects of neuroinflammation on whole neuron and synaptic compartments using a combination of MS, network analysis, confirmatory biochemical and ultrastructural assays and integrative approaches across our mouse-derived and existing human datasets.

**Results:** Ultrastructural and biochemical analyses of P2 fractions verified enrichment in synaptic elements, including synaptic vesicles and mitochondria. MS of biotinylated proteins from Camk2a-specific bulk brain homogenates (whole neuron) and P2 fractions (synaptosome) showed enrichment of >1000 proteins, consistent with neuron-specific biotinylation, also confirmed by immunofluorescence microscopy. Camk2a-specific synaptic proteome revealed molecular signatures related to mitochondrial function, synaptic transmission, protein translation. LPS-treated mice displayed body weight loss and neuroinflammation, characterized by glial activation, increased pro-inflammatory cytokine levels and upregulated expression of Alzheimer’s disease (AD)-related microglial genes. LPS-induced neuroinflammation exerted distinct effects on the synaptic proteome, including increased mitochondrial and reduced cytoskeletal-synaptic proteins, while suppressed synaptic MAPK signaling. Importantly, these changes were not observed at the whole neuron level, indicating unique vulnerability of the synapse to neuroinflammation. In line with synapse proteomic and signaling changes, LPS altered the ultrastructure of asymmetric synapses, suggesting dysregulation of excitatory neurotransmission. Co-expression network analysis of Camk2a neuronal proteins further resolved mitochondria- and synapse-specific protein modules, some of which were neuroinflammation-dependent. Neuroinflammation increased levels of a mitochondria-enriched module, and decreased levels of a pre-synaptic vesicle module, without impacting a post-synaptic membrane module. LPS-dependent mitochondrial and LPS-independent post-synaptic modules in mouse neurons mapped to post-mortem human AD brain proteomic modules which were decreased in cases with AD dementia and positively correlated to cognitive function, including pro-resilience markers for AD.

**Conclusion:** Our findings using native-state proteomics of Camk2a neurons combined with synaptosome enrichment identify proteome-level mechanisms of early synaptic vulnerability to neuroinflammation relevant to AD.

## BACKGROUND

Neurons span large distances in the nervous system, whereby neuronal soma and their projections (synapses/dendrites and axons), can function as molecularly distinct compartments. For instance, local translation of mRNA to proteins has been demonstrated to occur at the pre- and postsynaptic compartments (1, 2). Mitochondria are transported from the soma to the synapses to support high-energy demands for maintaining synaptic function and plasticity (3). Synaptic signaling mechanisms (e.g. MAPK and Akt/mTOR) are also known to regulate learning and memory via their critical roles in long-term potentiation (4, 5). As a result, it is likely that the synaptic compartment of neurons has differential vulnerabilities to mechanisms of neurological disease (6, 7). In neurodegenerative diseases, such as Alzheimer’s disease (AD), loss of proteins that localize to the synaptic compartment (e.g. cytoskeletal, scaffolding and synaptic vesicle proteins) is strongly correlated with cognitive decline. Therefore, not surprisingly, synaptic proteins have emerged as strong predictors of cognitive function and disease progression (8, 9).

In the early stages of AD, pathological processes (amyloid beta aggregation, tau tangle formation, and synaptic proteins changes) and cellular responses (microglia and astrocyte activation, impaired energy metabolism and mitochondrial dysfunction) occur prior to overt structural loss of synapses or neurodegeneration (10). Causal roles of pro-inflammatory mechanisms mediated by microglia, brain resident immune cells, have also been strongly suggested by identification of genetic risk factors in AD (e.g. APOE, TREM2, and C1Q) (11). Aging also increases neuroinflammatory responses in the brain, via activation of immune pathways including type 1 interferon signaling (12). Beyond AD, immune mechanisms primarily mediated by microglia and astrocytes, are implicated in several acute and chronic neuroinflammatory and non-AD neurodegenerative diseases (13). While mechanisms of neuroinflammation can vary across neurological diseases, pro-inflammatory responses can be reliably induced in mice using repeated systemic administration of lipopolysaccharide (LPS). This paradigm induces transient sickness response in mice (14), followed by reproducible microglial activation, upregulation of the complement system, interferon signaling and a unique gene expression profile (e.g. *Cst7, Clec7a, Itgax*) reminiscent of disease-associated microglia (DAM) in AD pathology (15–17). Importantly, the *in vivo* effects of LPS in this paradigm diverge from those observed *in vitro* (16, 18, 19), emphasizing the importance of maintaining the *in vivo* context. Neuroinflammation, aging and progressive AD pathology, ultimately lead to synapse loss, a key determinant of cognitive decline (20–26). Therefore, understanding how neuroinflammation impacts neurons and their synaptic compartments before overt neurodegeneration, is of critical importance in identifying the molecular basis of selective neuronal vulnerability and nominating targets for disease-modification (7, 27).

Bulk -omics analyses from post-mortem human brains can provide important insights into disease mechanisms (20–26), but are limited by sampling the end stage of disease, and might not accurately reflect the early stages or the disease trajectory. This bias becomes more important in diseases such as AD where the duration of the preclinical and the prodromal phases of disease pathogenesis can last years or even decades (28). Moreover, bulk-omics methods analyze a mixture of cells, masking the distinct protein expression profiles of individual cell types, with limited compartment-level resolution. Understanding molecular events occurring in the synapse, while retaining the native architecture of neurons in the brain, requires *in vivo* proteomic labeling using either bio-orthogonal (29) or proximity-labeling (for native-state proteomics) (30) or tagging of translational machinery (for translatomics) (31). These methods allow purification of tagged proteins or translated mRNAs for downstream ‘omics’ profiling.

One approach leverages the ability of biotin ligase TurboID to ubiquitously label lysine residues on proximate proteins (32). In our group, conditional TurboID expression in neurons has been achieved by crossing the Rosa26^TurboID^ mouse with an appropriate neuronal Cre-driver line, leading to TurboID expression only in neurons (33, 34). Selective biotinylated proteins can then be enriched from bulk brain homogenate, followed by proteomic characterization using label-free quantitative mass spectrometry (LFQ-MS). This approach is called cell type-specific in vivo biotinylation of proteins (CIBOP) and has been applied to neurons (Camk2a-CIBOP, PV interneuron-CIBOP) and astrocytes (Aldh1l1-CIBOP), yielding cell type-specific native-state proteomes (33–35). Recently, this approach successfully resolved early PV interneuron-specific proteomic changes in a mouse model of AD pathology, providing novel molecular signatures that could not be obtained in bulk tissue proteomics studies from the same animals (34). The ability of the CIBOP approach to label thousands of proteins in neurons across the soma, axons, dendrites and synapses, while retaining their native state in the brain, provides an opportunity to extend this approach for studying neuronal subcellular compartments. Biochemical subcellular fractionation methods to isolate synaptic compartments (synaptosomes) are a reliable tool to study synaptic proteomes in health and disease (36–40). While neuronal pre- and post-synaptic elements predominate in synaptic preparations, glial proteins are also highly abundant in these fractions making it difficult to completely delineate neuronal from glial proteins (41, 42). Labeling neuron-specific proteomes using CIBOP followed by synaptosome enrichment can distinguish neuronal from non-neuronal proteins, to resolve disease mechanisms specific to the synaptic compartment. We combined Camk2a-CIBOP with differential centrifugation to prepare enriched in pinched-off synapses (synaptosomes) to investigate the proteomic composition of the synaptic compartment of excitatory neurons in mouse brain. Using repeated systemic LPS to model neuroinflammatory stress, we contrasted the LPS-induced synaptosome proteomic alterations with those reflected in the whole-neuron proteome, complemented by synaptic ultrastructural analyses. We then applied a network-analysis approach to identify groups of highly co-expressed proteins, revealing increased mitochondrial proteins and decreased cytoskeletal/synaptic proteins and MAPK signaling pathways as unique effects of LPS on the excitatory synapse. These proteomic changes corresponded with subtle structural changes in pre-synapse size, synaptic vesicle density and PSD length. We also cross-referenced mouse-derived synaptic effects of neuroinflammation with post-mortem human and synaptosome proteomic networks, to identify molecular commonalities between synaptic effects of neuroinflammation and human AD pathogenesis.

## METHODS

### Animals

Experimental procedures were performed in accordance with the National Institutes of Health Guide for the Care and Use of Laboratory Animals. The protocol (PROTO-201700821) was reviewed and approved by the Emory University Institutional Animal Care and Use Committee. The experiments reported here are in accordance with the ARRIVE guidelines. Consistent with previous publications (33, 34) for our excitatory neuron-CIBOP approach, Rosa26^TurboID/wt^ (C57BL/6-Gt(ROSA)26Sortm1(birA)Srgj/J, JAX Strain #037890) mice were crossed with Camk2a-Cre^Ert2^ (B6;129S6-Tg(Camk2a-cre/ERT2)1Aibs/J, JAX Strain #012362) to obtain Rosa26^TurboID/wt^/Camk2a-Cre^Ert2^ (heterozygous for both) and their control littermates. All mice were given tamoxifen (75 mg/kg) intraperitoneally for 5 days starting at 8-9 weeks of age. After 3 weeks of recombination, mice were given water supplemented with biotin (37.5 mg/L) for 2 weeks. At the end of biotin supplementation, lipopolysaccharide (LPS, 500 µg/Kg, i.p., Sigma, L4391) was administered once daily for 4 consecutive days to induce acute neuroinflammation (16, 43), no mortality was observed during LPS administration, and euthanasia was performed 1 day after last dose of LPS. Male and female mice were used for the experiments with data collected from 4 mice per experimental condition. Animals were housed in the Department of Animal Resources at Emory University under a 12 h light/12 h dark cycle with *ad libitum* access to food and water and kept under environmentally controlled and pathogen-free conditions.

### Crude synaptosome preparation

Brain subcellular fractions were prepared as described previously (44, 45). Mice were deeply anesthetized using carbon dioxide and decapitated; their brains were quickly removed from the skull and dissected to exclude the cerebellum. All steps were performed either on ice or in a cold room. Each brain was gently homogenized with 16-strokes at 800 rpm in 10 volumes of Medium I (0.32 M sucrose, 5 mM HEPES pH 7.5, and 0.1 mM EDTA) containing protease and phosphatase inhibitors (ThermoFisher Scientific, 78446). The homogenate was first centrifuged at 1,000 x g for 10 min to give a pellet (P1) containing nuclear and cell debris. Supernatant (S1) was then centrifuged at 12,000 x g for 20 min to produce the crude synaptosome pellet (P2). Supernatant (S2) was discarded, the P2 pellet was resuspended in 500 µL Medium I and sonicated on ice until completely dissolved, and the resulting P2 fractions were stored at -80°C until used for downstream analysis.

### Transmission electron microscopy (TEM)

Fresh synaptosome-enriched P2 fraction pellets were fixed in a mixture of 2.0% paraformaldehyde and 2.5% glutaraldehyde in 0.1 M cacodylate buffer pH 7.4. The samples were next post-fixed with 1.0% osmium tetroxide and then stained en block in 2% aqueous uranyl acetate before being dehydrated through a series of graded ethanol concentrations up to 100% ethanol. Ethanol dehydration was followed by two changes in propylene oxide. Samples were then embedded in Eponate 12 resin and polymerized at 60°C for 48h. Ultrathin sections were cut on a Leica EM UC6 ultramicrotome, stained with uranyl acetate and lead citrate, and imaged in a Hitachi HT7700 TEM, operated at 80KV. For validation of enrichment, synaptosome images were collected at 5000x magnification.

For ultrastructural analysis, a separate cohort of wild-type (WT) mice treated with saline or LPS was deeply anesthetized and perfused with PBS followed by 4% paraformaldehyde (PFA). Whole brains were carefully dissected into two hemispheres and quickly fixed in 2.5% glutaraldehyde and 2% PFA in 0.1 M sodium cacodylate buffer (pH 7.4) for 1 h at room temperature and then kept overnight at 4°C. The next day, 150 µm-thick sagittal sections were cut on a vibratome, post-fixed for 1h on ice in the same fixative. The tissue was further post-fixed in 1% OsO4 and 0.8% potassium ferricyanide for 1 hour. Specimens were then en bloc stained with 2% aqueous uranyl acetate for 1 hour, dehydrated in a graded series of ethanol up to 100%, substituted with propylene oxide, and embedded in EMbed 812 resin (Electron Microscopy Sciences, Hartfield, PA). Sample blocks were polymerized in an oven at 60°C overnight. Ultrathin sections (60 nm) were cut using a Leica ultramicrotome (UC7) and post-stained with 2% uranyl acetate and lead citrate. The thin sections were examined with an FEI Tecnai transmission electron microscope at an 80 kV accelerating voltage, and digital images were acquired with an AMT NanoSprint15 MK2 camera (Advanced Microscopy Techniques, Woburn, MA. For the image analysis, the asymmetric synapse was defined as an electron-dense post-synaptic density area juxtaposed to a presynaptic terminal. Images used for analysis were acquired from layer 4-5 of the somatosensory cortex at 6800x magnification, and the analysis was performed while blind to the experimental conditions. Morphological features including presynaptic bouton size (area) and length of postsynaptic density (PSD) were manually delineated and measured using ImageJ, while the number of synaptic vesicles was manually counted and then divided by the presynaptic bouton area to estimate synaptic vesicle density.

### Sample preparation for protein-based analysis

Equal volumes of bulk brain homogenates and P2 fractions were diluted in 8 M urea lysis buffer (10mM TRIS, 100mM NaH2PO4, pH 8.5) with protease and phosphatase inhibitors (ThermoFisher Scientific, 78446), sonicated 3 times (Pulse 05 05; Amplitude 30%) on ice, and centrifuged at 18,000 x g for 10 min at 4°C. Supernatants were collected and total protein concentration was determined by BCA assay (ThermoFisher Scientific, 23225).

### Western blot (WB)

To confirm protein biotinylation, 20 µg of proteins from bulk brain homogenates and P2 fractions were separated on a 4-12% Bis-Tris gel (Invitrogen, NW04125BOX) at 80V and transferred to nitrocellulose membranes using the iBLOT mini stack system (Invitrogen, IB23002). Membranes were stained with Ponceau S (Sigma, P7170) for 2 min to confirm equal loading of gels. After washing with 0.1% TBS-Tween 20, the blots were blocked with blocking buffer (ThermoFisher Scientific, 37543) for 1h at room temperature (RT), probed with Streptavidin680 (Table 1) for 1h at RT and then imaged on the ChemiDoc Imaging System (Bio-Rad). To verify synaptic and mitochondrial protein enrichment, 4 µg of proteins from bulk brain homogenates and crude synaptosomal P2 fractions were resolved in a 4-20% Tris-Glycine gel (Invitrogen, XP04205BOX). After transfer, nitrocellulose membranes were probed with pre- and post-synaptic (SV2A, synaptophysin, synapsin-1, PSD95, complexin-1/2), mitochondrial (HSP60, SDHA, VDAC), cytosolic (α-tubulin) and nuclear (histone H3) markers (Table 1).

**Table 1.** List of primary and secondary antibodies used in this study.

A) Primary antibodies
| Target | Source | Cat # | Identifiers | Dilution | Application |
| --- | --- | --- | --- | --- | --- |
| PSD95 | Millipore | MAB1598 | RRID:AB_94278 | 1:500 | WB |
| Synaptophysin | Abcam | ab32127 | RRID:AB_2286949 | 1:20,000 | WB |
| Synapsin-1 | Cell Signaling | 5297 | RRID:AB_2616578 | 1:1,000 | WB |
| SV2A | Abcam | ab32942 | RRID:AB_778192 | 1:1,000 | WB |
| Complexin-1/2 | Cell Signaling | 28070 | RRID:AB_2798954 | 1:1,000 | WB |
| HSP60 | Cell Signaling | 12165 | RRID:AB_2636980 | 1:1,000 | WB |
| SDHA | Abcam | ab14715 | RRID:AB_301433 | 1:1,000 | WB |
| Histone H3 | Abcam | ab1791 | RRID:AB_302613 | 1:1,000 | WB |
| α-tubulin | Millipore | MAB1864 | RRID:AB_2210391 | 1:1,000 | WB |
| Iba1 | Abcam | ab178846 | RRID:AB_2636859 | 1:1,000 | IF |
| Iba1 | Synaptic Systems | HS-234 017 | RRID:AB_2891297 | 1:1,000 | IF |
| S100β | Synaptic Systems | 287 004 | RRID:AB_2620025 | 1:1,000 | IF |
| GFAP | Invitrogen | 13-0300 | RRID:AB_2532994 | 1:1,000 | WB, IF |
| NeuN | Synaptic Systems | 266 004 | RRID:AB_2619988 | 1:2,000 | IF |

B) Secondary antibodies
| Target | Source | Cat # | Identifiers | Dilution | Application |
| --- | --- | --- | --- | --- | --- |
| Streptavidin-DyLight 488 | Invitrogen | 21832 | N/A | 1:500 | IF |
| Streptavidin-Alexa Fluor 680 | Invitrogen | S32358 | N/A | 1:10,000 | WB |
| Donkey anti-rabbit-Alexa 680 | Invitrogen | A32734 | RRID:AB_2633283 | 1:10,000 | WB |
| Donkey anti-rabbit-Alexa 800 | Invitrogen | A32735 | RRID:AB_2633284 | 1:10,000 | WB |
| HRP anti-rabbit | Cell Signaling | 7074S | RRID:AB_2099233 | 1:10,000 | WB |
| Donkey anti-rat-IRDye CW 800 | LI-COR | 926-32219 | RRID:AB_1850025 | 1:10,000 | WB |
| Goat anti-mouse-Alexa 680 | Invitrogen | A32729 | RRID:AB_2633278 | 1:10,000 | WB |
| Goat anti-rabbit Alexa 555 | Invitrogen | A21428 | RRID:AB_2535849 | 1:500 | IF |
| Goat anti-guinea pig-Alexa 647 | Jackson Immuno Research | 106-605-003 | RRID:AB_2337446 | 1:500 | IF |
| Goat anti-rat-Alexa 488 | Invitrogen | A48262 | RRID:AB_2896330 | 1:500 | IF |
| Donkey anti-rabbit-Alexa 647 | Invitrogen | A32795 | RRID:AB_2762835 | 1:500 | IF |

IF indicates immunofluorescence; WB indicates western blot

### Affinity purification of biotinylated proteins

Following our previously optimized protocol (33, 34), biotinylated proteins were captured by streptavidin magnetic beads (ThermoFisher Scientific, 88817) incubating 41.5 µL beads per 500 µg of protein in 500 µL RIPA buffer (50 mM Tris, 150 mM NaCl, 0.1% SDS, 0.5% sodium deoxycholate, 1% Triton X-100) for 1h at 4°C with rotation. After incubation, beads were then washed sequentially at RT 2x with RIPA buffer for 8 min/each, 1x with 1 M KCl for 8 min, 1x with 0.1 M Na2CO3, 1x with 2 M urea in 10mM Tris-HCl (pH 8.0) for 10 sec, 2x with RIPA buffer for 8 min/each and 4x PBS. Finally, beads were then recovered on a magnetic rack, PBS was removed completely and further diluted in 100 µL PBS. For quality control studies by WB and silver staining, 10% of beads were eluted by heating the beads in 30 µL of 2X protein loading buffer (Bio-Rad, 1610737) supplemented with 2 mM biotin + 20 mM dithiothreitol (DTT) at 95°C for 10 min. Subsequently, 10 μL of eluate was run on a gel and probed with Streptavidin680 while 20 μL of eluate was run on a separate gel and Silver stained (ThermoFisher Scientific, 24612). The remaining enriched biotinylated proteins were stored at -20°C until on-bead digestion.

### Protein digestion, peptide clean up and label-free quantitative mass spectrometry (LFQ-MS)

On-bead digestion of proteins from streptavidin-enriched pulldown samples and digestion of inputs (bulk brain homogenates and P2 fractions) were processed similarly to our previous CIBOP studies (33, 34). In brief, the remaining 90% of streptavidin beads were washed with PBS and then resuspended in 225 μL of 50 mM ammonium bicarbonate (ABC, NH4HCO3). About 1 mM DTT was added to reduce samples for 30 min on a shaker (800 rpm) at RT. About 5 mM iodoacetamide was added to alkylate cysteines for 30 min on a rotor at RT, protected from light. Proteins were digested overnight with 0.5 μg of lysyl endopeptidase (Lys-C; Wako, 127-06621) on a shaker (800 rpm) at RT. Next day, proteins were further digested by 1 μg trypsin (ThermoFisher Scientific, 90058) overnight on a shaker (800 rpm) at RT. Samples were acidified to 1% formic acid and 0.1% TFA, desalted using HLB columns (Waters, 186008055), and dried using cold vacuum centrifugation (SpeedVac Vacuum Concentrator).

To prepare inputs via in-solution digestion, 50 µg of protein from bulk brain homogenates and P2 fractions were reduced with 5 mM DTT at RT for 30 min and alkylated by 10 mM IAA in the dark for 30 min. Samples were then digested overnight with 1 µg of Lys-C at RT on a shaker. Samples were further diluted 7-fold with 50 mM ABC and digested overnight with 1 µg of trypsin at RT on a shaker. The peptide solutions were acidified to a final concentration of 1% FA and 0.1% TFA, desalted with HLB columns (Waters, 186008055), and dried down in a vacuum centrifuge (SpeedVac Vacuum Concentrator). Dried peptides were resuspended in loading buffer (0.1% FA and 0.03% TFA in water), then analyzed by liquid chromatography and MS (Q-Exactive Plus, Thermo, data dependent acquisition mode). Approximately 1ug (inputs) or 10% (pulldowns) was loaded onto a Water’s CSH column (150um x 15 cm with 1.7um CSH beads) with a Dionex RSLCNano and eluted over a 60 minute gradient. Buffer A consisted of water and 0.1% formic acid and buffer B was comprised of 80% acetonitrile and 0.1% formic acid. Spectra were collected on an Orbitrap Eclipse mass spectrometer. Survey scans were collected at 120,000 resolution, mass range from 400 to 1600, automatic gain control (AGC) set at 4 x e5 and maximum injection time to 246ms. Tandem scans were collected at 30,000 resolution, isolation width at 1.6 mass over charge, 35% collision energy, normalized AGC at 100% and 54 ms max injection time. The mass spectrometer was set to collect at top speed for cycles of 3 seconds. Only ions with a charge state from 2 to 5 were chosen for fragmentation and dynamic exclusion time was set to 30 seconds with a +/- 10 ppm mass tolerance.

### Proteomic pipeline of inputs and streptavidin-enriched pulldown samples

In bulk brain homogenates and P2 fraction lysates (inputs), analysis yielded 2,540 protein identifications across 32 samples following 25% filtration in Perseus. For Camk2a whole-neuron homogenate and P2 fraction pulldowns, label-free quantification (LFQ) intensities were processed through a standardized Perseus workflow. Potential contaminants were excluded, and the data were log2-transformed. Protein recovery varied depending on the filtration threshold, with 1,447 proteins retained at 50%, 1,741 at 25%, and 1,985 at 10%. Missing values were imputed from a normal distribution to correct data sparsity, and the dataset was subjected to quantile normalization to minimize technical variance across samples. Quality control was performed by boxplot visualization of LFQ distributions before and after normalization, as well as principal component analysis (PCA) to assess clustering and variance structure. The processed matrix was then exported for downstream analyses. Differential abundance testing was conducted on both bulk input and pulldown datasets using log2-transformed, normalized intensities. Pairwise comparisons employed two-tailed unpaired t-tests with equal variance assumed, while multi-group comparisons used one-way ANOVA with post-hoc Tukey HSD. Both raw and FDR-corrected p-values were calculated, although unadjusted p-values together with fold-enrichment estimates were emphasized to increase analytical stringency. Differentially expressed proteins (DEPs) were visualized using volcano and heatmap plots, applying a significance threshold of *p* < 0.05 and |log2FC| > 0.

### Synaptic Gene Ontologies and Annotations

Synaptic proteins were annotated using the SynGO Database (46) [available from (47)], which provides curated classifications of synaptic genes by cellular component and biological process. Protein identifications were mapped to SynGO entries via HGNC-approved gene symbols and categorized into presynaptic, postsynaptic, and synaptic vesicle compartments, as well as functional processes. To expand this analysis, annotations from SynaptopathyDB (48), a consensus set of 3,437 mammalian synapse proteins from presynaptic and postsynaptic compartments, were also incorporated. Synaptic protein sets identified in the P2-pulldown were intersected with MitoCarta3.0 (49), that contains 1,140 mouse proteins with strong evidence of mitochondrial localization [available from (50)], to identify mitochondria-associated synaptic proteins.

### Gene Ontology (GO) Analysis

GO enrichment analysis was carried out using the clusterProfiler v.3.21 and enrichR v.3.4 packages, with the mouse genome annotation as the reference set and the experimentally identified proteome as the background. Enrichment was assessed across the three principal GO domains Biological Process (BP), Molecular Function (MF), and Cellular Component (CC). Results were visualized in ggplot2, with terms ranked by -log10 adjusted p-value, and the most significantly enriched categories within each domain highlighted for downstream interpretation.

### Gene Set Variation Analysis (GSVA)

GSVA was performed using the GSVA R package (51) with a custom GMT file, applying the z-score algorithm to compute pathway enrichment scores across samples. Pathway score correlations were evaluated by Pearson correlation with BH-adjusted p-values and displayed as heat maps. Robustness was evaluated by 1,000 permutations, with empirical p-values derived from the proportion of permuted mean differences exceeding the observed mean difference.

### Ingenuity Pathway Analysis

Pathway and network analyses were performed using QIAGEN Ingenuity Pathway Analysis (IPA) software v.24.0.1 [available from (52)] to identify canonical pathways, upstream regulators, and molecular networks altered by LPS. Analyses were conducted separately for bulk brain homogenates and P2 fraction lysates (inputs) and for Camk2a whole-neuron homogenate and P2 fraction pulldowns, followed by comparative evaluation across all datasets. Pathway activity was inferred from IPA z-scores, with values > 0 considered activated and values < 0 considered inhibited. Significance thresholds were set at p < 0.05 (–log10 p > 1.3).

### Mitochondrial abundance-based normalization

Mitochondrial abundance-based normalization was used to determine whether changes in specific mitochondrial proteins were due to LPS and not to changes in mitochondrial mass. Mitochondrial mass was estimated using a sum of protein abundances of all mitochondrial proteins in our data. Inputs and pulldown proteomes were overlapped with the mitochondrial reference protein set derived from the MitoCarta3.0 (49). The sum abundance of mitochondrial proteins (n=256) in the Camk2a TurboID-pulldown samples that mapped to the MitoCarta3.0 protein list, was used to scale proteomic data from each sample to its mitochondrial content. Differential enrichment analysis for LPS effect were then repeated.

### Cognitive Resilience Analysis

Resilience-associated PWAS enrichment was evaluated for Camk2a neuron-enriched proteins (homogenate and P2 fractions pulldowns) using a permutation-based framework (22) implemented with MAGMA.SPA [available from (53)]. Reference data were obtained from the PWAS of cognitive trajectory by Yu *et al*. (22), which tested 8,356 proteins for associations with longitudinal cognitive change. Proteins with FDR < 0.2 were retained, with positively associated genes defined as pro-resilience (n = 300) and negatively associated as anti-resilience (n = 216).

### Weighted Gene Co-expression Network Analysis (WGCNA)

A scale-free co-expression network was constructed using the WGCNA R package v.1.73 (54). The optimal soft-thresholding power (β) was selected with pick Soft Threshold to approximate scale-free topology. Pearson correlations were used to generate signed adjacency matrices, and Topological Overlap Measure (TOM) dissimilarities were computed. Proteins were hierarchically clustered, and modules were identified with cut-tree Hybrid, with stability confirmed across deep split parameters.

### Module preservation across Human TMT proteomics

Module preservation between mouse Camk2a and published human TMT proteomic data was evaluated using the module preservation function (500 permutations; minimum module size =15). Following preservation analysis, eigengene-based connectivity (kME) was calculated, representing the correlation between each gene’s expression profile and its corresponding module eigengene. Module eigengenes were computed to identify the genes within each co-expression module, and module membership was defined as the correlation between individual gene expression profiles and the respective module eigengene. Module membership and eigengene connectivity (kME) were quantified, and the significance of module overlap between datasets was assessed using Fisher’s Exact Test (*p*<0.05). Pearson correlations between each protein and each module eigen protein were performed.

### Functional Annotation and Pathway Enrichment of Predicted Modules

To evaluate the biological significance of the identified modules, functional annotation and pathway enrichment analyses were performed. Gene Ontology (GO) enrichment was conducted with the clusterProfiler and enrichR packages, using the complete protein dataset as background and the Mouse Genome Database [available from (55)] as reference. Enrichment was assessed across the three GO domains Biological Process (BP), Molecular Function (MF), and Cellular Component (CC) to identify pathways and functional categories associated with each module. GO terms with p < 0.05 were considered significantly enriched.

### Protein-Protein Interaction (PPI) Network and Hub Gene Identification

Protein-protein Interaction network was constructed from differentially expressed proteins (DEPs) to examine predicted protein interactions using Cystoscope software v.3.2.0 [available from (56)]. In the network, nodes represent genes and edges represent the interactions between the nodes.

### Differential localization of ribosomal proteins in Camk2a homogenate and P2 fraction pulldowns

Given local protein synthesis occurs in neuronal compartments, including pre- and postsynaptic compartments (1, 57), we utilized the HUGO [Human Genome Organization, available from (58)] ribosomal protein sets to identify and intersect ribosomal proteins detected in the Camk2a homogenate and P2 fraction pulldowns.

### Human synaptosome data FET analysis

Human synaptosome proteomic data from Carlyle *et al*. (59) were used to test enrichment of up- and downregulated proteins within Camk2a-centered WGCNA modules. Fisher’s Exact Test was applied using all quantified proteins as the background, and significance was reported as –log₁₀(p-values) with thresholds (p < 0.05, *p < 0.01, **p < 0.001).

### Immunofluorescent staining and image analysis

Free-floating (40 µm-thick) sagittal sections were incubated for 30 min with 10 mM sodium citrate buffer pH 6 at 65°C for antigen retrieval, blocked with 5% bovine serum albumin (BSA) (Jackson ImmunoResearch, 001-000-162) in 0.3% PBS-Triton X-100 (PBS-T, Sigma, T8787) for 1 h at RT, and then incubated overnight at 4°C with primary antibodies (Table 1A) GFAP and S100β (astrocyte markers), Iba1 (microglial marker), or NeuN (neuronal marker) in 1% BSA. The next day, sections were washed 3x with PBS-T and then incubated with corresponding secondary antibodies (Table 1B) for 1 h at RT in the dark, followed by nuclear staining with DAPI [1µg/mL] (Roche, 10236276001) for 2 min. After washing 2x with PBS, sections were mounted with ProLong Diamond antifade mounting medium (Invitrogen, P36961); once dried, slides were stored at 4°C in the dark. Fluorescence images of somatosensory cortex layer 5 and hippocampal CA1 region consistently across all samples were acquired at 20x magnification using the Keyence BZ-X810 microscope. A threshold level for each glial marker was established to detect differences between groups, which was used for all images within a given marker, using the NIH ImageJ Fiji software. Percentage area occupied by each glial marker, denoted here as % area covered, indicates the area above the threshold level. For assessing neuronal density with NeuN, manual cell counts were obtained for each group. For each marker, a mean ± standard error of the mean (SEM) was calculated for each group (n = 4–6). The data were analyzed by unpaired t-test for comparison between saline- and LPS-cohorts.

### Quantification of cytokines and signaling phospho-protein levels

Multiplexed Luminex immunoassays were used to measure cytokines (Millipore, MCYTMAG-70K-PX32), phospho-proteins in the MAPK (Millipore, 48-660MAG) and PI3/Akt/mTOR pathways (Millipore, 48-612MAG) in bulk-brain homogenates and P2 fractions as previously described (33, 34). Briefly, cytokines were analyzed using the standard Luminex assay, first magnetic beads were incubated with sample, followed by biotinylated detection antibodies, and then streptavidin-phycoerythrin. The magnetic beads uniquely identify each analyte and the streptavidin-phycoerythrin provides relative analyte quantity. On the other hand, MAPK and mTOR signaling were analyzed using the adapted assay, which takes advantage of the Camk2a neuron-specific biotinylated proteome, omitting the biotinylated detection antibody step, to directly quantify biotinylated neuron-derived proteins. A linear range analysis was conducted to identify the range of total protein loaded that corresponded to a linear signal readout from the Luminex instrument for each analyte. Based on this, 1.25 μg of total protein was loaded for each sample for the cytokine, PI3K/Akt/mTOR, and MAPK assays, respectively. Luminex assays were read on a MAGPIX system. Luminex data were z-scored and discriminant partial least squares regression analysis (D-PLSR) was performed using the opls function in the ropls package v.1.36.0 in R, after the data were z-scored. Standard deviation of latent variable one (LV1) loading for each protein was determined using a leave-one our cross validation (LOOCV) approach. In this approach the D-PLSR projection was recalculated 100 times with a random sample omitted each time. The output of D-PLSR was rotated to best capture group-wise differences in LV1.

### Enzyme-linked immunosorbent assay (ELISA)

For plasma preparation, after decapitation, trunk blood samples were collected in Eppendorf tubes with EDTA and centrifuged at 1,000 x g for 5 min at 4°C. Plasma was collected, diluted [1:25] in assay buffer, and analyzed using a commercial ELISA kit (UmanDiagnostics, 20-8002) for the neurofilament light chain (NfL), a biomarker of axonal injury in neurodegenerative diseases, as per manufacturer’s instructions.

### Acute isolation of CD11b+ myeloid cells from mouse brain

For gene expression analysis, a separate cohort of WT mice treated with PBS or LPS (500 µg/Kg, i.p., once daily for 4 consecutive days) were deeply anesthetized, perfused with cold PBS, and quickly decapitated. Brains were removed, and hemispheres separated along the midline, a piece of left hemisphere was snap-frozen for bulk brain tissue RNA-Seq studies. Microglia cells were isolated from the right hemisphere by Percoll gradient centrifugation as previously described (15), with minor modifications. Briefly, brain tissue was mechanically dissociated on a 40 µm cell strainer over a 50 mL conical tube, suspended in 30 mL PBS and centrifuged for 5 min at 800 x g at 4°C. Cell pellets were then resuspended in 6 mL 35% stock isotonic Percoll solution (90% Percoll + 10% 10X HBSS) and transferred to 15mL conical tubes. The cells were centrifuged for 25 min at 800 x g with a speed of 2 for deceleration at 4°C. The floating myelin layer was removed by aspiration. After washing with PBS, cells were resuspended in 200 μL PBS containing transcription inhibitors (10 μM Triptolide, 27.1 μg/mL Anisomycin, 5 μg/mL Actinomycin, 1:100 Ribolock). Forty μL CD11b+ magnetic beads (Miltenyi Biotec, 130-093-636) were added to the cell suspension and incubated protected from light for 15 min. Following incubation, cells were resuspended in PBS and passed through LC columns. Following complete flow through, cells were centrifuged at 800 x g for 5 min at 4 °C to remove PBS and snap-frozen. From frozen bulk brain tissue and CD11b+ microglia cells, total RNA was isolated with TRIzol (Invitrogen, 15596018) and after assessing RNA quality/purity, the cDNA library was prepared. We analyzed RNAseq data from brain and from CD11b+ microglia in contrast for LPS vs WT using DEseq2 method and visualized differential expression genes using volcano plots.

### Statistical analysis and figure preparation

All analyses of proteomic and RNAseq data were done in R. Visualization of results used figures generated from R and edited in Adobe Illustrator. In the proteomics study, a sample was detected as an outlier and then excluded from the MS analysis. Graphs and figures summarizing *in vivo* studies and biochemical studies were prepared using GraphPad Prism v.10.2.3, BioRender.com and Adobe Illustrator. Additional statistical considerations specific to each figure are also provided in the figure legends.

## RESULTS

### Native-state proteomics of excitatory neuron-specific crude synaptosomes

Bulk tissue homogenates prepared from fresh mouse brains were fractionated by differential centrifugation to obtain crude synaptosomal P2 fractions (**Fig. 1A**). TEM confirmed enrichment of pinched-off synaptic terminals within the P2 fractions, containing synaptic vesicles and mitochondria (**Fig. 1B**). Immunofluorescence microscopy of Camk2a-CIBOP brain tissue confirmed cell type-specific biotin labeling of excitatory neurons, including soma and neuronal projections (**Fig. 1C**), with localization of biotin signal within β3 tubulin-positive neurons but absent in non-neuronal cells. Western blot (WB) analysis of bulk brain homogenates and P2 fraction lysates (inputs) using streptavidin to detect biotinylated proteins, showed robust biotinylation in Camk2a-CIBOP mice compared with minimal endogenous biotinylation in negative controls (**Fig. 1D**, *top panel*). Like inputs, WB of streptavidin-enriched Camk2a whole-neuron samples (homogenate pulldowns) and Camk2a synaptosome samples (P2 fraction pulldowns) showed mirroring patterns of strong protein biotinylation in Camk2a-CIBOP mice compared to negative controls (**Fig. 1D**, *bottom panel*). The silver stain of eluates also confirmed enrichment of biotinylated proteins from Camk2a-CIBOP mice (**Supplementary Fig. S1A**). WB further confirmed that P2 fractions were enriched in synaptic (SV2A, synaptophysin, synapsin-1, PSD95, complexin-1/2) and mitochondrial (HSP60, SDHA, VDAC) markers, while depleted in cytosolic (α-tubulin) and nuclear (histone H3) markers (**Fig. 1E**). Based on ultrastructural and biochemical validation of P2 synaptosome preparations and neuron-specific protein biotinylation, we then performed label-free quantitative mass spectrometry (LFQ-MS) on input (bulk brain homogenate and P2 synaptosomal fraction referred henceforth as ‘homogenate-input’ and ‘P2-input’) as well as enriched biotinylated proteins obtained from streptavidin-pulldown samples (referred to as ‘homogenate-pulldown’ and ‘P2-pulldown’ fractions), and quantified 2,540 proteins and 1,741 proteins, respectively. Principal-component analysis (PCA) of Camk2a-specific proteomes revealed separated clustering of Camk2a-CIBOP samples and negative controls (**Supplementary Fig. S1B**). The first and second principal components represent variation caused by mouse line (PC1= Camk2a-CIBOP mice vs non-CIBOP negative controls; 25% variance) and subcellular fractionation (PC2= Bulk brain homogenates vs crude synaptosomal P2 fractions; 11% variance).

**Figure 1.**
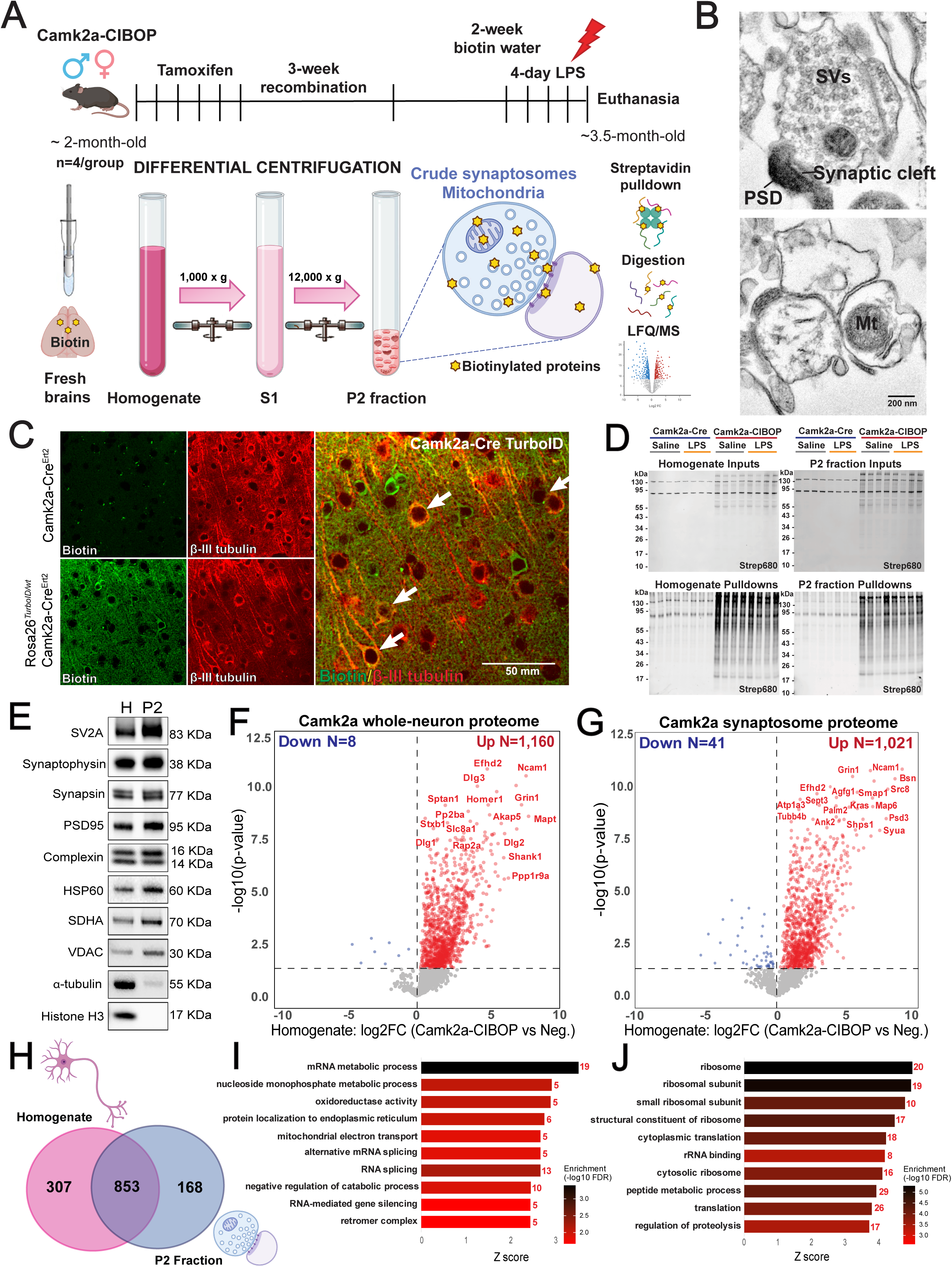
Native-state proteomic profiling of Camk2a excitatory neuron-specific crude synaptosomes. **(A)** Experimental outline to achieve native-state proteomics of Camk2a excitatory neuron-specific crude synaptosomes. **(B)** Representative electron micrographs of P2 fractions confirmed that crude synaptosomes have sealed plasma membrane, filled with synaptic vesicles and intact mitochondria, synaptic cleft and postsynaptic terminals with electron-dense post-synaptic density. SVs, synaptic vehicles; PSD, post-synaptic density; Mt, mitochondria. **(C)** Representative immunofluorescence images of cortex from non-CIBOP negative controls (Camk2a-Cre^Ert2^) and Camk2a-CIBOP (*Rosa26^TurboID/wt^*/Camk2a-Cre^Ert2^) mice showing biotinylation (green streptavidin Alexa488) in relation to cell body (red neuron-specific β-III Tubulin). **(D)** Western blots of input and streptavidin affinity pulldown samples confirm strong protein biotinylation in Camk2a-CIBOP mice as compared to limited biotinylation in non-CIBOP negative controls. **(E)** Representative blots of the subcellular fractionation show enrichment of synaptic (SV2A, synaptophysin, synapsin-1, PSD95, complexin-1/2) and mitochondrial markers (HSP60, SDHA, VDAC) in crude synaptosome fractions (P2) compared to bulk brain homogenates (H), while depletion of cytosolic (α-tubulin) and nuclear (histone H3) markers, 4 μg of protein was loaded for each fraction. Volcano plots showing differentially enriched proteins in **(F)** bulk brain homogenate and **(G)** P2 fraction pulldowns comparing Camk2a-CIBOP mice vs negative controls. Red dots represent proteins biotinylated in Camk2a whole-neuron and Camk2a synaptosome proteomes (unpaired two-tailed t-test *p*<0.05). **(H)** Venn diagram showing the number of identified proteins and overlap between Camk2a whole-neuron and Camk2a synaptosome proteomes. Examples of unique enriched proteins in **(I)** Top enriched pathways for the 307 proteins identified exclusively in the Camk2a whole-neuron proteome. **(J)** Top enriched pathways for the 168 proteins identified exclusively in the Camk2a synaptosome proteome. Protein abundance is plotted as log2-transformed and normalized intensity values (*n*=4 biologically independent samples). Also see Additional files 1 and 7 for related analyses and datasets, and Additional file 12 for Supplementary Fig. S1. Image was created using BioRender.com.

First, to characterize the Camk2a whole-neuronal proteome, we compared bulk brain homogenate-pulldowns from Camk2a-CIBOP samples with non-CIBOP control pulldown samples and identified 1,160 proteins that were biotinylated in Camk2a CIBOP samples. These included proteins involved in neurotransmission (Grin1), synaptic signaling (Ncam1, Dlg1-3, Shank1, Homer1), and cytoskeletal dynamics (Efhd2, Palm2, Mapt, Rap2a, Sptan1) (**Fig. 1F**), consistent with a neuronal proteome. Similarly, to determine the Camk2a-specific synaptosome proteome, we compared P2-pulldown proteomes from Camk2a-CIBOP mice with all non-CIBOP P2-pulldown controls and identified 1,021 biotinylated proteins. These included proteins involved in synaptic transmission (Ncam1, Bsn, Ank2, Syua, Psd3) and mitochondria energy metabolism (Tom20, Ndua4, Atp5i) (**Fig. 1G**), again consistent with a neuronal proteome. Interestingly, beyond canonical markers of excitatory neurons, we also identified 168 proteins exclusively enriched in the Camk2a-specific synaptosome proteome that were not present in the whole-neuronal proteome, such as Nars1, Prkca, Rpl28, Shank2, Arf5, Tom20, Mark2 and Vdac1 (**Fig. 1H, Additional file 1**). Moreover, contrasting with the homogenate-pulldown proteome (**Fig. 1I**), the top pathways for the 168 proteins identified exclusively in the P2-pulldown proteome included ribosome, rRNA binding, and translation (**Fig. 1J**), indicating that our Camk2a-specific synaptosome proteome also captured the protein synthesis machinery for local translation at neuronal synapses. These data demonstrate that our Camk2a-CIBOP approach combined with subcellular fractionation reveal a unique synaptic compartment-specific proteomic signature in excitatory neurons otherwise overseen in the whole-neuronal proteome.

### Verification of synaptic and mitochondrial enrichment by subcellular fractionation and CIBOP

The homogenate-input proteome reflects the bulk brain tissue proteome while P2-input proteome captures the proteome of crude synaptosomes enriched in synaptic and mitochondrial elements, some of which are neuronal while others are glia-derived. By imposing CIBOP and enrichment of biotinylated proteins to homogenate and P2 fraction samples, we expect homogenate-pulldown proteomes to be enriched in somatodendritic/cytoskeletal and nuclear neuronal proteins while P2-pulldown proteomes will be enriched in neuron-specific synaptic and mitochondrial proteins as mitochondria account for 30% of the volume of a glutamatergic synapse (60). To verify these predicted patterns at the proteomic level, we compared P2-input with homogenate-input (**Fig. 2A**), as well as P2-pulldown with homogenate-pulldown proteomes (**Fig. 2B**). We identified 581 proteins increased in P2-input, while 540 proteins were increased in homogenate-input proteomes (**Fig. 2A**). The P2-input proteome was enriched in mitochondrial proteins (Slc25a5, Sdha, Gls, Vdac1), while de-enriched in nuclear (Septin8, Anp32a, Actc1) and mRNA splicing-related proteins (Srsf2, Snrpd1, Eef2) (**Fig. 2C**). To determine proteomic composition of Camk2a-specific synaptosomes, we compared P2-pulldown with homogenate-pulldown and identified 233 proteins increased in P2-pulldown and 221 increased in homogenate-pulldown (**Fig. 2B**). The P2-pulldown proteome was enriched in mitochondrial (Ndufs2, Sucla2, Slc25a4), ribonucleotide binding (Gnao1, Pccb, Opa1) and synaptic terms (Flot2, Gpr158, Grin1) while the homogenate-pulldown proteome was enriched in myelin sheath, axon and cell body/cytoskeletal terms (**Fig. 2D**). Furthermore, we found 127 unique proteins most-highly abundant in the Camk2a-specific synaptosomal P2-pulldown proteome (**Fig. 2E**). These include mitochondrial, synaptic transmission, as well as ribosomal proteins potentially involved in local protein translation at synaptic compartments (**Fig. 2F, Supplementary Fig. S1C**). Using the synaptic protein database SynGO (46), the 233 synaptic proteins enriched in the P2-pulldown proteome included pre- and post-synaptic proteins (**Fig. 2G**), as well as plasma membrane channels such as Lrrc8a, Gria2, Grin2a, Grin2b, Grin1, and Kcnma1. Given that mitochondrial proteins are underrepresented in SynGO, SynaptopathyDB database (48) was used for further analysis of P2-pulldown proteome and then intersected with the MitoCarta3.0 inventory (49) to identify mitochondria-associated synaptic proteins. We found 24 mitochondrial proteins allocated in the pre-synaptic compartment, 40 mitochondrial proteins allocated in the post-synaptic compartment, and 21 mitochondrial proteins not specifically allocated to pre- or post-synaptic compartments (**Additional file 2**). Collectively, these results validate our Camk2a-CIBOP approach combined with subcellular fractionation for native-state proteomic studies of whole neurons and more specifically, their synaptic compartment.

**Figure 2.**
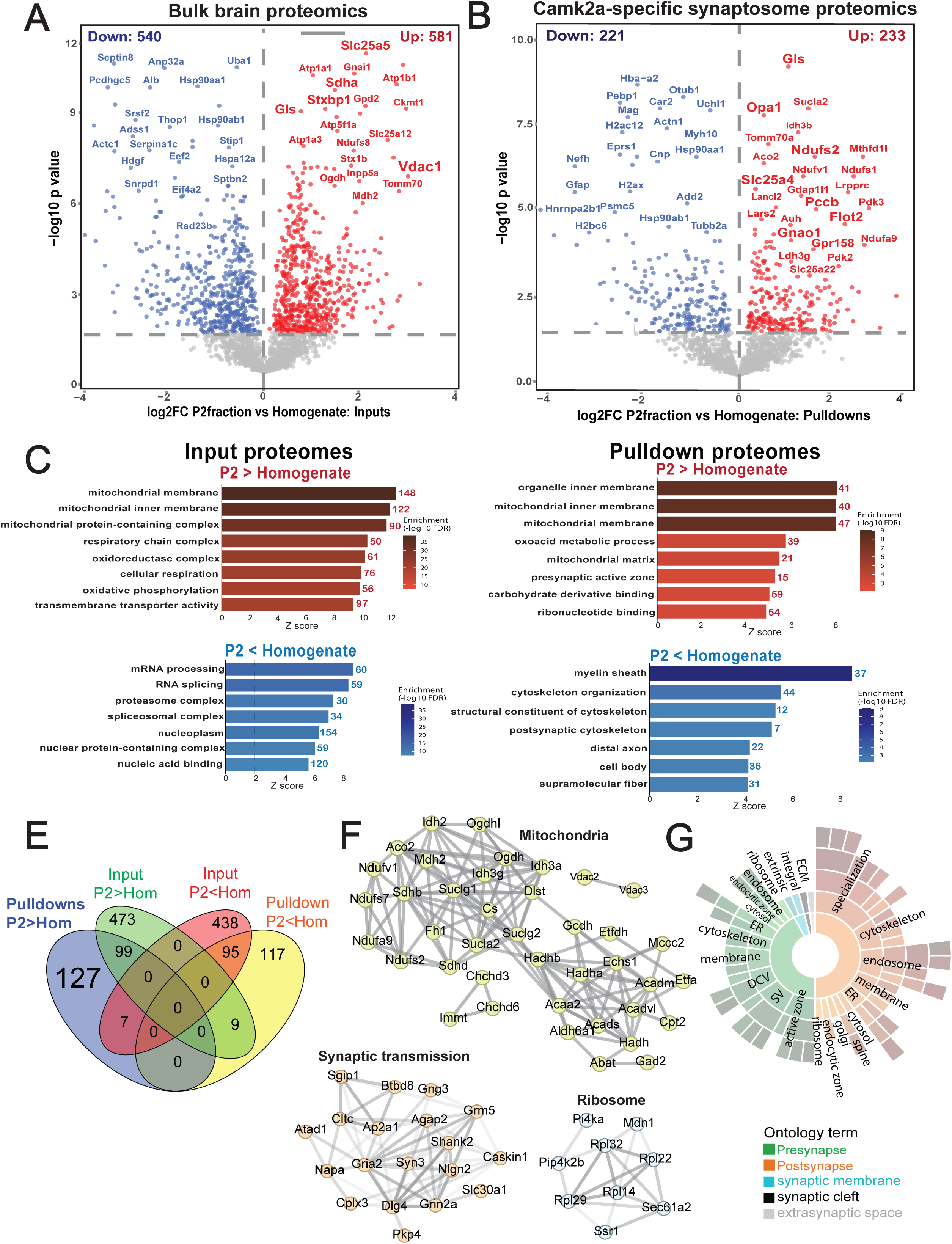
Volcano plots show differentially enriched proteins in **(A)** bulk brain samples (P2-input vs homogenate-input) and **(B)** Camk2a-neuron specific samples (P2-pulldown vs homogenate-pulldown). **(C)** Gene Ontology (GO) analyses of brain synaptosome (P2-input) proteomes showing enrichment of mitochondrial membrane, cellular respiration, and transporter activity proteins, while depletion in mRNA processing, proteasome complex, and nucleic acid binding proteins. **(D)** GO analysis of Camk2a-specific synaptosome (P2-pulldown) proteomes showed enrichment of mitochondrial membrane, ribonucleotide binding, and presynaptic active zone, while depletion of myelin sheath, cytoskeleton organization and cell body proteins. **(E)** Venn diagram of the unique and shared proteins that were identified from P2 fraction inputs and pulldowns. **(F)** STRING network of the 127 unique increased proteins identified in Camk2a-specific synaptosomes showing enrichment in mitochondria, synaptic transmission, and ribosomes. **(G)** SynGO cellular component analysis of the 233 increased proteins in Camk2a-specific synaptosomes reveals labeling of proteins in the pre- and post-synaptic compartments. Also see Additional file 2 for related analyses and datasets.

### Modeling neuroinflammation relevant to neurodegeneration, using repeated LPS administration

Following validation of excitatory synaptosome-specific proteomics using Camk2a-CIBOP and subcellular fractionation, we aimed to identify synapse-specific effects of neuroinflammation. To elicit neuroinflammation in the mouse brain, we used the systemic LPS model in which four daily intraperitoneal doses cause sickness behavior, weight loss and robust microglial activation in the brain (14, 43, 61). Despite LPS does not directly entering the brain, it mediates neuroinflammation via indirect pathways (61, 62). As expected, both CIBOP and non-CIBOP mice treated with LPS experienced weight loss from day 2 to day 4 of LPS treatment compared to saline controls (**Fig. 3A**). Microglia and astrocyte activation was observed in the hippocampal CA1 region (**Fig. 3B, Supplementary Fig. S2A and D**) and the somatosensory cortex (**Fig. 3B, Supplementary Fig. S2B and E**) of LPS-treated mice, evidenced by higher percentage area covered and increased numbers of Iba1+ and S100β+ cells, respectively. Despite robust microglial activation and reactive astrogliosis, LPS-treatment did not impact plasma NfL levels (**Fig. 3C**), nor the density of NeuN-positive neurons (**Supplementary Fig. S2C and F**), indicating lack of overt axonal injury nor neurodegeneration in this model at 24 h after LPS administration.

**Figure 3.**
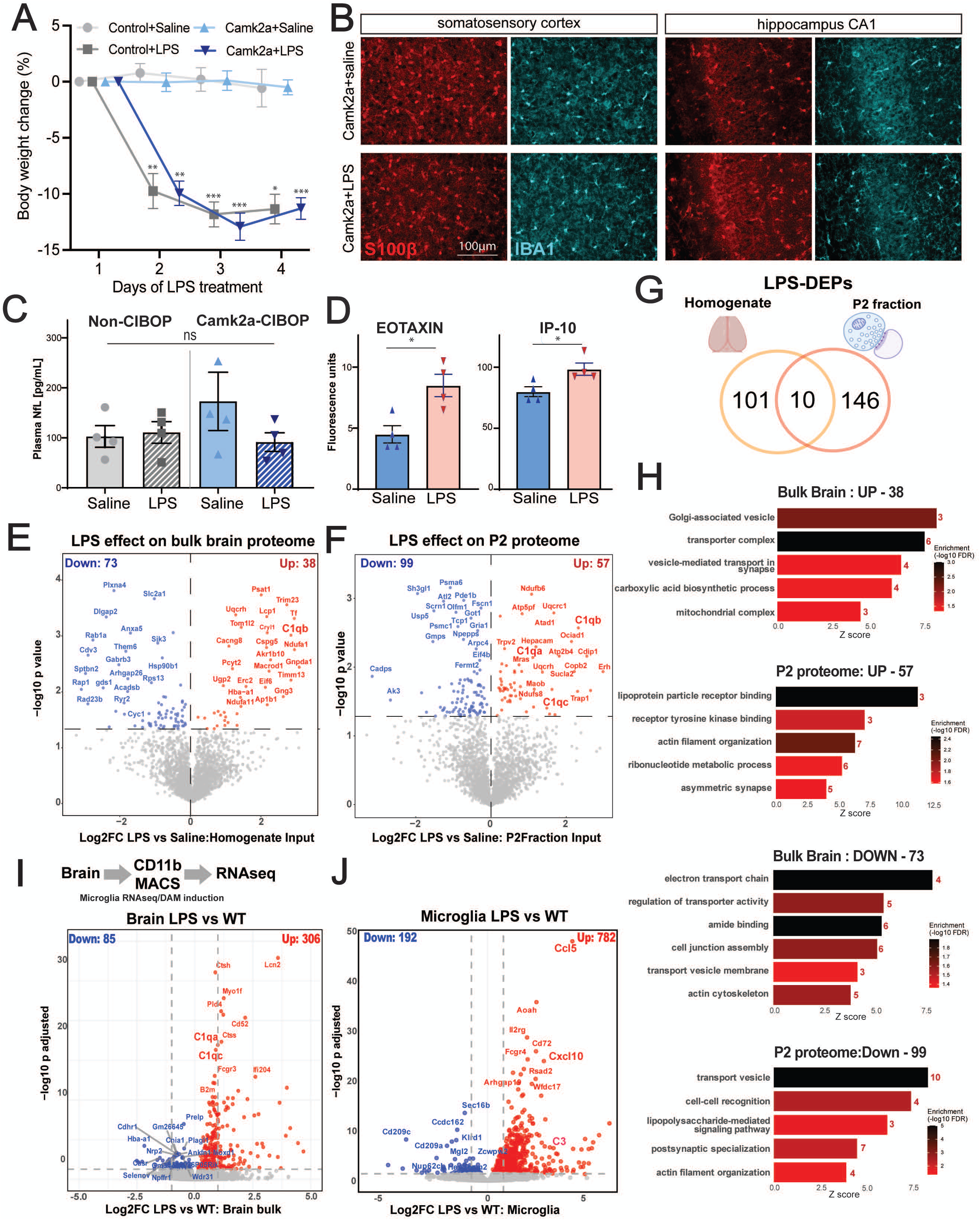
Global effects of systemic LPS. **(A)** LPS-treated mice exhibited a significant reduction in body weight. Two-way repeated measures ANOVA [F(9, 36)=14.37, *p*<0.0001] followed by Tukey’s multiple comparison test (\**p*<0.05, \*\**p*<0.01, \*\*\**p*<0.001). **(B)** Representative immunofluorescence images showing the distribution of Iba1^+^ cells (microglia in cyan), and S100β^+^ cells (astrocytes in red) in the somatosensory cortex and hippocampal CA1 region and from Camk2a-CIBOP mice treated with LPS compared to saline controls. Systemic LPS administration induced microglial activation and astrocyte reactivity. **(C)** LPS-treated mice exhibited similar NF-L plasma levels than saline controls; non-CIBOP+saline vs non-CIBOP+LPS, 102.8 ± 21.7 pg/mL vs 110.7 ± 21.6 pg/mL; Camk2a-CIBOP+saline vs Camk2a-CIBOP+LPS, 172.9 ± 58.4 pg/mL vs 91.4 ± 18.5 pg/mL. Data are expressed as mean ± SEM. One-way ANOVA [F(3, 12)=1.135, p=0.3741], ns= no significant. **(D)** Selected chemokine, cytokine, and growth factor levels were measured in bulk brain homogenates using a standard multiplexed Luminex assay. Upregulation of Eotaxin and IP-10 expression were the most relevant protein changes in LPS-treated mice. Data are expressed as mean ± SEM. Unpaired two-tailed t-test [Eotaxin t=3.464, df=6, p=0.0134; IP-10 t=2.846, df=6, p=0.0293] (\**p*<0.05). Volcano plots showing the effects of LPS on **(E)** bulk brain proteomes and on **(F)** P2 synaptosomal fraction proteomes. **(G)** Venn diagram of the unique and shared proteins that were induced by LPS in bulk brain homogenates and P2 fractions. **(H)** Results of GO enrichment analysis of bulk brain homogenates (LPS homogenate-input vs Saline homogenate-input) and P2 fractions (LPS P2-input vs Saline P2-input). Selected GO terms from each of the three GO groups (biological process, cellular component, and molecular function) are shown. Volcano plots of differentially expressed gene transcripts comparing LPS-treated to saline-treated WT mice in **(H)** brain and **(I)** microglia. Also see Additional files 3, 8, 9 for related analyses and datasets, and Additional file 12 for Supplementary Figs. S2 and S3.

Pro-inflammatory cytokine levels (Eotaxin and IP-10/Cxcl10) were significantly increased in LPS-treated mice as compared to saline-treated mice (**Fig. 3D**). In bulk brain homogenates, LPS treatment increased levels of 38 proteins and decreased levels of 73 proteins (**Fig. 3E-G**). In the bulk brain (homogenate-input) proteome, LPS-increased levels of proteins involved in intracellular protein transport, carboxylic acid biosynthetic and mitochondrial respiration. Conversely, LPS reduced levels of proteins involved in aerobic electron transport chain, cytoskeleton organization, and postsynaptic specialization (**Fig. 3H**). In P2 fractions, LPS increased levels of 57 proteins and decreased levels of 99 proteins (**Fig. 3F-G**). In the P2 (P2-input) proteome, LPS treatment increased proteins involved in mitochondrial respiration, actin filament organization, and asymmetric synapse. Conversely, LPS reduced levels of proteins involved in vesicle transport and synaptic vesicle-related functions (**Fig. 3H**). Importantly, only 10 proteins overlapped in between conditions (**Fig. 3G**), suggesting that the effect of neuroinflammation on bulk brain homogenates and synapse/mitochondria-enriched P2 fractions are molecularly distinct.

We then asked whether the inflammatory stress imposed by the LPS-treatment paradigm has similarities to neuroinflammatory mechanisms observed in AD. In independent experiments, we isolated CD11b+ MACS-purified brain myeloid cells (>95% microglia) from LPS-treated and saline-treated WT mice and performed RNAseq to identify transcriptomic effects of LPS on brain and microglia. In bulk brain tissues, LPS treatment upregulated 306 genes (*Saa3, Adgre1, Lcn2, Ctss*) and downregulated 85 genes (*Plagl1, Prelp, Nrp2, Chia1*) (**Fig. 3I**). In microglia, LPS treatment resulted in 782 upregulated genes (*Ccl5, Aoah, Cd72, Cxcl10, Lyz2*) and 192 downregulated genes (*Ccdc162, Cd209c, Nup62c, Sec16b*) (**Fig. 3J**). The larger magnitude of effect of LPS on microglial transcriptomes than bulk brain transcriptomes indicates that microglia may be primary responders to LPS-mediated neuroinflammatory stress in this model. LPS-upregulated genes in microglia included pro-inflammatory (*Ccl5, Cxcl10, Il1b*) and several canonical DAM genes (*Cd72, CD69, Stat1 and Treml2*), while downregulated genes included several homeostatic markers (*Ccnd1, Fmn2, Fbxl13*). These results suggest that LPS-induced neuroinflammation recapitulates several aspects of microglial activation observed in models of neurodegeneration such as AD. For instance, neuroinflammatory mediators, such as complement C3 and C1q, have been implicated in microglia-mediated synapse loss in AD (63–65). Since we observed LPS-induced increased C1q levels in bulk brain homogenate and P2 fraction proteomes, we compared C1q protein induction in our LPS studies with bulk brain proteomic data from WT and 5xFAD mice, euthanized at 2 different ages 6-and 11-months of age for early and late amyloid-β pathology, respectively. C1qa, C1qb, C1qc proteins were consistently increased across LPS-treated mice and 5xFAD mice (**Supplementary Fig. S3**). In summary, we observed that systemic LPS-induced neuroinflammation induces robust microglial activation with several similarities to microglial phenotypes observed in amyloid-β pathology models. The AD-relevant microglial responses due to LPS treatment, are also associated with some distinct effects on the P2 fraction involving mitochondrial, synaptic and cytoskeletal protein alterations, as well as complement-related mechanisms that are of relevance to AD. Since the crude synaptosomal P2 fraction contains neuronal and non-neuronal elements, enrichment of biotinylated proteins from Camk2a-CIBOP animals could more precisely delineate neuroinflammation-induced effects on the synaptic compartment of excitatory neurons.

### LPS-neuroinflammation differentially impacts the proteome of Camk2a neuron synaptic compartments and mitochondria, with corresponding ultrastructural changes

We examined the effects of LPS-neuroinflammation on Camk2a-specific whole-neuron proteomes, and more specifically, on its synaptic compartment. LFQ-MS of biotinylated proteins obtained from homogenate-pulldown (whole-neuron proteome) (**Fig. 4A**) and from P2-pulldown (synaptosome proteome) (**Fig. 4B**) samples identified 59 LPS-induced DEPs in whole-neuron proteomes and 55 DEPs in the synaptosome proteome, of which only 2 DEPs overlapped (Anp32a, Ctsd) (**Fig. 4C**). Particularly, in synaptosomal P2 fractions, LPS increased levels of mitochondrial proteins (Pdhb, Ndufc2, Uqcrh), Rac/Rab GTPases (Rac1, Rab35), and decreased levels of cytoskeleton (Map4), synaptic vesicle (Rab3gap1, Snap47), translation (Eif6), and calcium-signaling (Calb2) proteins (**Fig. 4B**). GSVA of LPS effects on the whole-neuron and synaptosome proteomes further identified unique effects of neuroinflammation on synapses which were not observed at the level of whole-neuron proteomes (**Fig. 4D**). For instance, LPS whole-neuron proteomes showed upregulation of detoxification and oxidoreductase activity, while a reduced neuron-synapse, somatodendritic compartment, and cytoskeleton organization pathways. In synaptosomal P2 fraction proteomes, LPS increased proteins involved in mitochondrial acyltransferase and metabolic activity, including purine containing compound metabolic processes, but decreased proteins involved in nucleoside triphosphate regulator activity. Conserved effects of LPS included increased mitochondrial aerobic respiratory chain-related proteins at both whole-neuronal and synaptosome proteomic levels (**Fig. 4D**). Supporting GSVA data, IPA of whole-neuron and synaptosome proteome changes revealed a similar pattern of LPS-induced alterations in mitochondria-associated pathways, highlighting that in P2-input proteome, LPS increased oxidative phosphorylation and respiratory electron transport, while reduced mitochondrial dysfunction. Similarly, in P2-pulldown proteome, LPS increased respiratory electron transport (**Supplementary Fig. S4C-H**). Together, these findings suggest that neuroinflammatory stress induces mitochondrial compensatory bioenergetic signatures prominent at the synapses.

**Figure 4.**
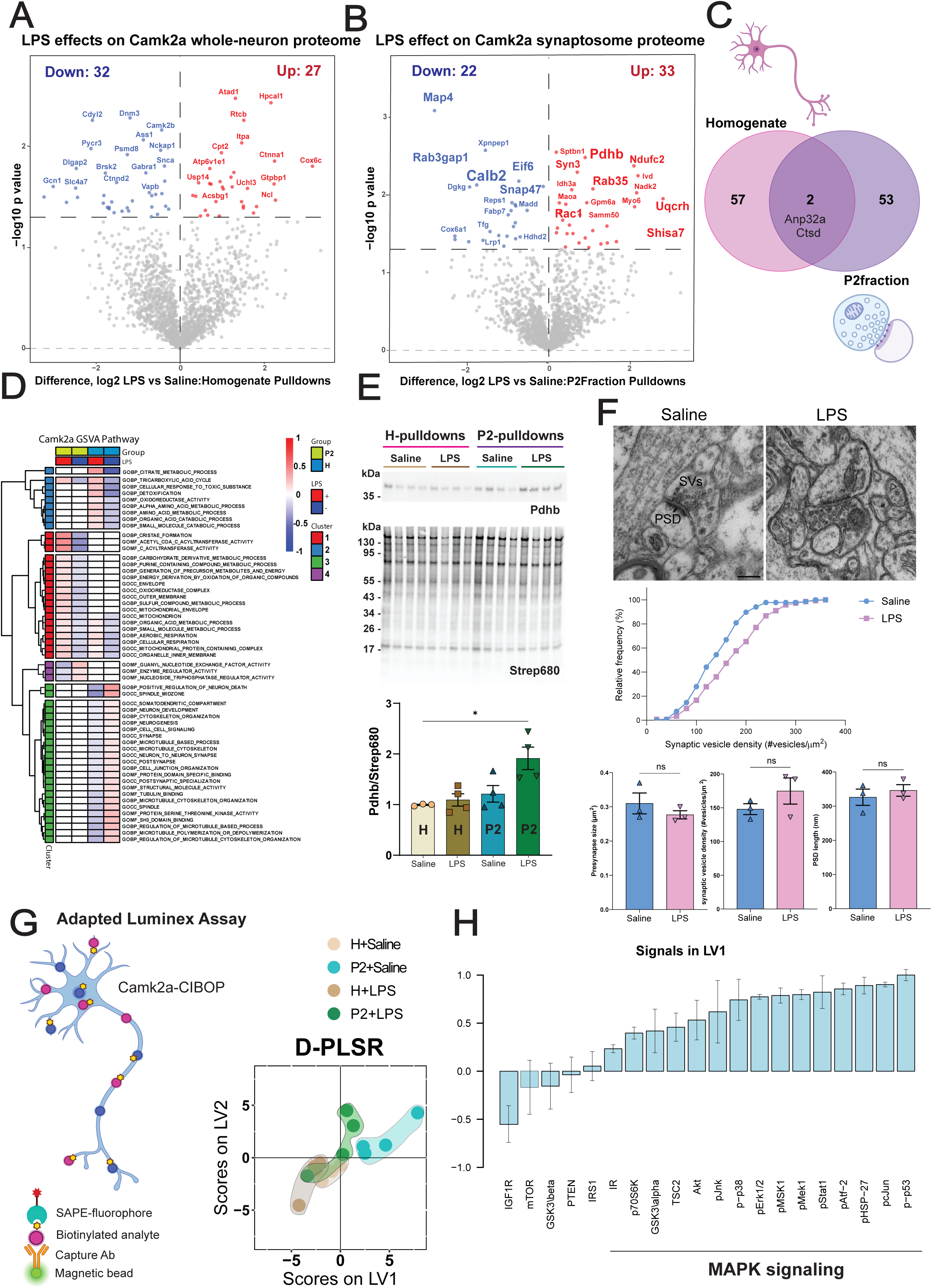
LPS induced proteomic, ultrastructural and signaling changes in excitatory synapses. Volcano plots showing the effects of LPS on **(A)** Camk2a whole-neuron proteome and **(B)** Camk2a synaptosome proteome. **(C)** Venn diagram of the unique and shared proteins that were induced by LPS in Camk2a-CIBOP homogenates and P2 fractions. **(D)** GSVA reveals differential effects of LPS on pathway enrichment in homogenates and P2 fractions. **(E)** WB verification of increased Pdhb protein levels in Camk2-CIBOP P2-pulldowns of LPS-treated mice as compared to H-pulldowns and saline controls (*n* = 3‒4). Data shown as mean ± SEM. One-way ANOVA [F(3,11)=6.628, p=0.0081], followed by Tukey’s multiple comparison test (\**p*<0.05). **(F**, *top panel*) Representative electron micrographs showing asymmetric synapses in the somatosensory cortex of WT mice treated with saline or LPS. **(F**, bottom *panel*) Cumulative distribution of synaptic vesicle density LPS-treated mice displayed an increased synaptic vesicle density [Kolmogorov–Smirnov (K-S) test, D=0.2443, p=0.0005]. Saline *n*=136 and LPS *n*=144 asymmetric synapses from three WT mice for each treatment. When parameters were averaged per animal, no significant differences were found in the quantification of the presynapse size [K-S test, D=0.3333, *p*>0.9999], synaptic vesicle density [K-S test, D=0.6667, p=0.6], or PSD length [K-S test, D=0.3333, *p*>0.9999], ns=no significant. The scale bar represents 200 nm. Data are presented as mean ± SEM. **(G**, left) Cartoon representation of adapted Luminex assay to measure the levels of Camk2a neuron-derived phospho-proteins belonging to MAPK and Akt/mTOR signaling pathways from bulk brain homogenates and P2 fractions. **(G**, right) D-PLSR analysis was used to cluster the samples based on treatment and sample type. **(H)** Individual analyte loadings on LV1 Positive values indicate the analyte is associated with positive scores on LV1 and vice versa for negative loadings. Error bars indicate standard deviation of loadings from leave-one out cross validation. Also see Additional files 4, 10, 11 for related analyses and datasets, and Additional file 12 for Supplementary Figs. S4 and S5.

A very important advantage of combining CIBOP with synaptosome enrichment for proteomic analysis is the identification and quantification of low-abundance proteins located at the synapses (e.g. Pdhb, Ndufc2, Uqcrh, Syn3, Calb2, Snap47, Map4), with no change detectable at whole-neuron homogenate level. Given the functional relevance of mitochondrial proteins uniquely increased by LPS in Camk2a synaptosome proteomes, we validated the increase of Pdhb (subunit of pyruvate dehydrogenase crucial mitochondrial enzyme for energy production) by WB (**Fig. 4E**). Using Camk2a-CIBOP homogenate-pulldowns and P2-pulldowns, Pdhb was significantly increased in P2-pulldowns but not homogenate-pulldowns from LPS-treated mice (**Fig. 4E** and **Supplementary Fig. S4A**). No effect of LPS in Pdhb protein level was observed in the bulk brain tissue proteome (homogenate-input) nor synaptosome proteomes (P2-input) from Camk2a-CIBOP mice (**Supplementary Fig. S4B**). Interestingly, we did not observe changes in mitochondrial function (Rptor) or dynamics regulators (Dnm1l, Mfn2, Opa1) in P2-pulldown proteomes from LPS-treated mice. To determine whether changes observed in mitochondrial proteins in neurons due to LPS-induced neuroinflammation can be explained by changes in total mitochondrial mass versus other mechanisms, we estimated mitochondrial mass using a sum of protein abundances of all mitochondrial proteins in our data. We found that LPS increased mitochondrial mass comparing LPS-treated group to saline controls within inputs but only in P2 fractions, while no significant differences were found within pulldowns, in either homogenates or P2 fractions. We also normalized our homogenate and P2 pulldown data for mitochondrial abundances, and observed the same DEPs (including Pdhb) as a result of LPS effect. While not directly conclusive, our results suggest that the observed effects of LPS-induced neuroinflammation on neuronal synaptic mitochondrial proteins are not likely due to alterations in mitochondrial mass in each compartment (**Supplementary Fig. S5**).

To further understand whether the LPS-induced alterations of synaptic proteins observed in Camk2a synaptosome proteomes may also impact synaptic structure, we examined the ultrastructural features of asymmetric (predominantly excitatory) synapses in the somatosensory cortex using TEM (**Fig. 4F**, *top panel*). We compared several parameters across LPS and non-LPS groups at the level of individual synapses in the TEM micrographs and the cumulative distribution curve revealed that synaptic vesicle density was significantly increased, while the pre-synaptic bouton size and PSD length did not change in the LPS group, as compared to saline-treated mice (**Fig. 4F**, *middle panel* and **Supplementary Fig. S4I-J**). However, no statistically significant group differences were found when parameters were averaged per animal (**Fig. 4F**, *bottom panels*). Together, the ultrastructural study results showing increased synaptic vesicle density under neuroinflammatory stress, suggests that LPS may enhance excitatory synaptic transmission, consistent with alterations of synaptic proteins (e.g. decreased Rab3gap1 and Snap47, increased Syn3) in Camk2a synaptosome proteomes from LPS-treated mice.

### Camk2a neuron synaptic compartment exhibits differential regulation of MAPK signaling in response to neuroinflammatory stress

Several pathways including mitogen activated protein kinase (MAPK) and Akt/mTOR signaling are important regulators of synaptic function and plasticity (4, 5). MAPK regulates synaptic activities, in fact several synaptic proteins have been identified as MAPK substates, including scaffolding proteins (PSD95), cadherin-associated proteins, potassium channels, and metabotropic glutamate receptors. The phosphorylation of these proteins results in their trafficking and synaptic delivery and thus determines the strength and efficacy of excitatory synapses. Activation of these pathways, triggered by an upstream activating signal, leads to sequential phosphorylation of activity-regulating residues on protein kinases, ultimately regulating key transcriptional factors which impact down-stream gene regulation involved in synaptic plasticity (5). We have recently leveraged Camk2a-CIBOP to label neuron-specific signaling proteins, along with an adapted Luminex approach to directly quantify neuronally derived signaling phospho-proteins from brain tissue lysates (**Fig. 4G**) (33, 66). We applied this approach to measure phospho-proteins (MAPK and Akt/mTOR pathways) derived from Camk2a neurons, using bulk brain homogenate-input (whole-neuron) and P2-input (synaptosome) lysates and examined the effect of LPS-neuroinflammation on these signaling pathways. At basal conditions (without LPS), D-PLSR analysis revealed that MAPK pathway activation was higher in the synaptic compartment (exemplified by a positive signal of LV1 for pJnk, pMek1, p-p38 and pErk1/2) as compared to the whole-neuronal level (**Fig. 4G-H**). We also observed that the effect of LPS-neuroinflammation on these signaling pathways was most apparent in the P2 synaptosome proteome as opposed to the whole-neuronal homogenates (**Fig. 4 G-H**), emphasizing that the impact of LPS on these signaling pathways was preferential to the synaptic compartment. Specifically, MAPK signaling was suppressed by LPS-neuroinflammation, while Akt/mTOR signaling was not impacted (**Fig. 4G-H**).

### Co-expression network analysis identifies LPS-dependent and -independent mitochondrial and synaptic modules

To complement differential enrichment analyses, we examined our Camk2a whole-neuron proteomes (homogenate-pulldown) and Camk2a synaptosome proteomes (P2-pulldown) using WGCNA. This dimensionality-reduction strategy allows us to understand changes occurring at the level of functional groups of co-expressed proteins (modules) rather than individual protein levels. WGCNA applied to 1,741 proteins identified 14 co-expression modules (M1-M14, module size range 57-204) displayed as a module heatmap and Eigen protein network dendrogram (**Fig. 5A-B**). Whole-neuron proteomes displayed higher expression in protein modules M1 (Turquoise) enriched in somatodendritic proteins (Mbp, Tubb, Ptpa) and M4 (Yellow) enriched in spliceosome and nuclear protein terms (Map2, Nefl, Nefh) (**Fig. 5C** and **E**). In contrast, Camk2a P2 synaptosome proteomes showed higher levels in M2 (Blue) module enriched in mitochondria (Ndufs2, Opa1, Gls), M9 (Magenta) module enriched in pre-synaptic vesicle and GABAergic transmission (Eif4b, Sncg, Cttn), M6 (Red) module enriched in post-synaptic membrane and glutamatergic transmission (Dlg4, Grin1, Grin2a, Gria3), and M5 (Green) module enriched in mitochondria (Ndfs7, Slc25a11, Ndufb4) and lipid metabolism terms (Snd1, Ppt1) (**Fig. 5D** and **E**). While none of the whole-neuron modules were impacted by LPS (**Fig. 5C**), modules M2 and M5 (mitochondria), and M9 (synaptic vesicle and GABAergic transmission) showed LPS effects, where modules M2 and M5 were increased, but M9 decreased in P2 fraction-pulldowns from LPS-treated mice (**Fig. 5D**). This observed pattern suggests that neuroinflammation exerts a stronger effect on the synaptic compartment as compared to the somatodendritic compartment of neurons. Furthermore, within the synaptic compartment, neuroinflammation impacts mitochondria, synaptic vesicles and GABAergic neurotransmission while glutamatergic transmission and the post-synaptic compartment may be relatively spared.

**Figure 5.**
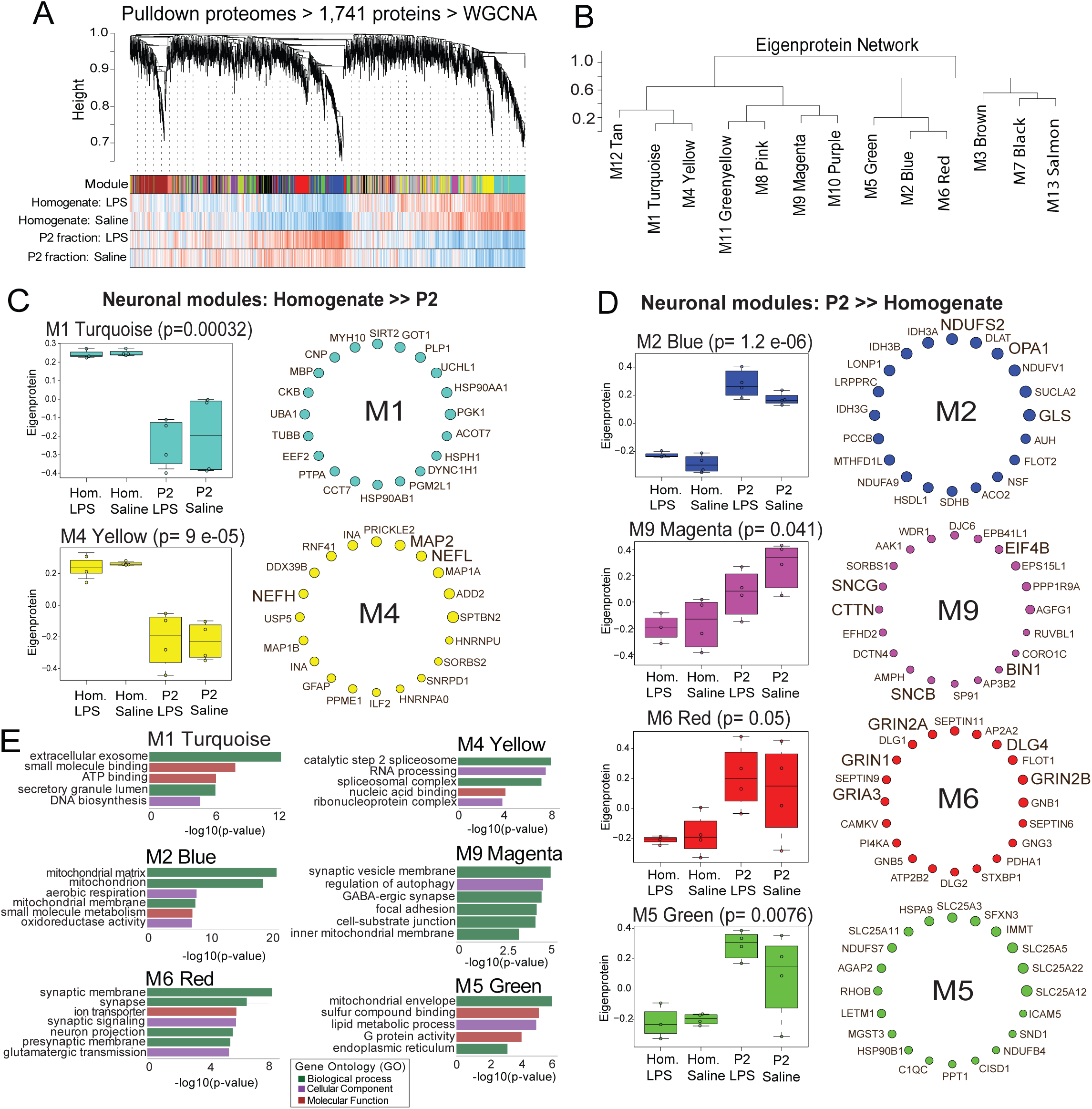
Co-expression network analysis of excitatory neuronal proteins. **(A)** Hierarchical clustering dendrogram of 1,741 proteins identified from homogenate-pulldown and P2-pulldown proteomics. Modules were identified by dynamic tree cut, and the corresponding module colors are displayed in the heatmap. Sample traits such as experimental group (Homogenate vs. P2 fraction) and treatment (LPS vs Saline) are shown beneath the dendrogram. **(B)** Eigenprotein network dendrogram of co-expression modules. **(C)** Modules enriched in Camk2a-CIBOP homogenate-pulldowns. Left Boxplots showing significantly higher eigengene expression in Homogenate versus P2 fractions for modules M1 (turquoise; p = 0.00032) and M4 (yellow; p = 0.05). Right Circular network visualizations of hub proteins within each module, annotated with gene symbols. These modules were enriched in somatodendritic and nuclear proteins. **(D)** Modules enriched in Camk2a-CIBOP P2 fractions. Left Boxplots showing significantly higher eigengene expression in P2 fractions versus Homogenate for M2 (blue; p = 0.09), M9 (magenta; p = 0.041), M6 (red; p = 0.05), and M5 (green; p = 0.0079). Right Circular network visualizations of hub proteins within each module, annotated with gene symbols. These modules were enriched in mitochondrial and synaptic proteins. **(E)** Gene ontology (GO) enrichment analysis of neuronal modules. Top GO terms (biological processes) are shown for M1 Turquoise, M4 Yellow, M2 Blue, M9 Magenta, M6 Red, and M5 Green. Bar plots display -log10(p-value) for enrichment, color-coded by GO category. Also see Additional file 5 for related analyses and datasets.

### Synapse-enriched protein modules are preserved in a proteomic network derived from post-mortem human brain

To assess the human translational relevance of mouse neuronal and synaptic co-expression networks, we assessed how protein modules identified in our dataset from Camk2a-CIBOP mice (homogenate-pulldown and P2-pulldown) are conserved in a proteomic network identified using human post-mortem bulk brain proteomes (21). By using TMT-labeled mass spectrometry (TMT-MS), this reference human dataset identified over 8,000 proteins from 488 human post-mortem brain tissues (21), and the WGCNA-derived network comprised 42 modules (**Fig. 6A**). Several mouse modules demonstrated strong preservation in the human network, particularly M2 (mitochondria), M6 (synapse-glutamatergic), and M9 (synaptic vesicle and GABAergic transmission). Notably, these three mouse modules showed higher levels in the synapse-enriched compartment as compared to whole neuronal proteome (**Fig. 5D**).

**Figure 6.**
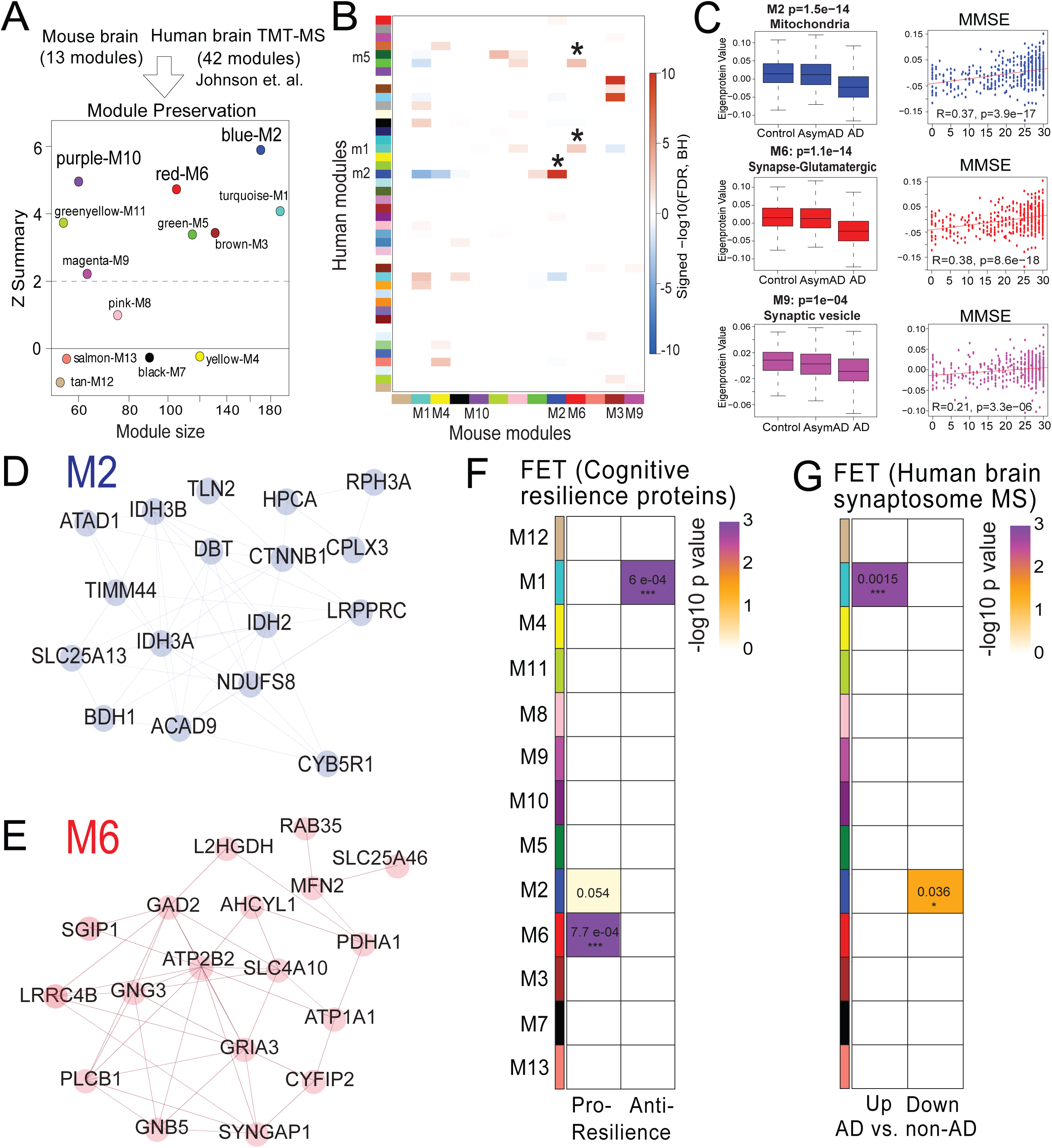
Integration of mouse and human data revealed molecular signatures associated with cognitive resilience. **(A)** Module preservation analysis between mouse and human TMT-MS proteomes. The Z summary preservation score (y-axis) is plotted against module size (x-axis) for 13 mouse modules tested for preservation in the human dataset (42 modules, ∼8,000 proteins) from PWAS data (21). Mouse modules M2 Blue and M6 Red show strong preservation in the human network. **(B)** Cross-species module overlaps heatmap. Rows represent human modules; columns represent mouse modules. The color intensity reflects the degree of overlap based on protein membership, with red indicating greater overlap. Mouse modules M2 and M6 demonstrated the greatest overlap with human modules. **(C)** Synthetic eigenprotein module analysis. Left panels show box plots comparing eigenprotein expression levels between AD cases and controls across modules M2, M6 and M9. The right panels display scatterplots showing a significant correlation between module eigenprotein expression and cognitive performance, as measured by Mini-Mental State Examination (MMSE) scores [M2 Blue R=0.37, p=3.9e−17, M6 Red R=0.38, p=8.6e−18, M9 Magenta R=0.21, p=3.3e−06]. STRING network for pro-resilience proteins from modules **(D)** M2 and **(E)** M6. Overlap of mouse modules [M1 Turquoise (somatodendritic, LPS-independent in Homogenate), M2 Blue (mitochondria, LPS-increased in P2 fraction), and M6 Red (synapse-glutamatergic, LPS-increased in P2 fraction)] with human protein modules by Fisher’s Exact Test (FET). The color scale ranges from 0 (white, no enrichment) to 3 (purple, FET P value ≤ 1e-3). Y-axis Mouse WGCNA modules; X-axis Human proteomics modules from the **(F)** cognitive resilience study (22) and the **(G)** human synaptosome study (59). Module M6 (p=0.0007) show strong enrichment and module M2 (p=0.054) a trend for pro-resilience, while module M1 (p=6e-04) shows strong enrichment for anti-resilience proteins. Also see Additional file 6 for related analyses and datasets.

Focusing on these mitochondria/synapse-enriched modules preserved across species, we then examined how these modules changed across Control (no pathology), Asymptomatic AD (AsymAD, AD pathology without cognitive symptoms), and Alzheimer’s Disease (AD pathology with cognitive symptoms) groups in the human proteome (**Fig. 6B**). Interestingly, the mouse M2 ‘mitochondria/metabolism’ module (matched to human M2 Blue module) which increased with LPS, was significantly decreased in human AD compared to control groups (**Fig. 6C**, left). The mouse M6 ‘post-synaptic/excitatory’ module (matched to human M1 Turquoise and M5 Green modules), and the mouse M9 ‘inhibitory/synaptic vesicle’ module (matched to human M39 module), both displayed a significant decrease in AD compared to the control group (**Fig. 6C**, left). Moreover, module eigen proteins for these conserved neuronal modules (mouse M2, M6, M9) were significantly and positively correlated with cognitive performance, as measured by Mini-Mental State Examination (MMSE) scores (**Fig. 6C**, right). Based on these patterns, we infer that the effects of LPS-neuroinflammation on neuronal mitochondria and synapses most likely represent early neuronal events in AD pathology, which ultimately decrease due to overt neurodegeneration. Whether the early increase in neuronal mitochondrial protein levels due to LPS-neuroinflammation represents early dysfunction or compensation to maintain cellular homeostasis, remains unclear.

### Cognitive resilience proteins are enriched in neuroinflammation-related mitochondrial and synaptic protein modules

To determine whether LPS-induced neuroinflammation impacts cognitive resilience mechanisms in Camk2a neurons and synapses, we performed FET enrichment analysis using lists of pro-resilience and anti-resilience proteins previously defined from PWAS of the human dorsolateral prefrontal cortex (22). We found that M2 (mitochondria) and M6 (synapse-glutamatergic) modules were significantly enriched in pro-resilience proteins while the M1 Turquoise (somatodendritic neuronal) module was enriched in anti-resilience proteins (**Fig. 6D**). Key pro-resilience proteins from M2 module include Rph3a, Ndufs8, Atad1, Idh3b, Hpca, and Timm44 (**Fig. 6E**), which are involved in mitochondrial import, redox homeostasis, and tricarboxylic acid cycle integrity-functions that maintain neuronal energy metabolism and potentially prevent bioenergetic collapse in AD. Pro-resilience proteins from M6 include Gria3, Gng3, Rab35, Plcb1 and Syngap1 (**Fig. 6F**), are implicated in excitatory neurotransmission, synaptic plasticity, and calcium regulation, supporting a model in which preserved glutamatergic synaptic function promotes resistance to cognitive decline. Next, we integrated our synapse-enriched neuron-specific proteomics analysis with a recent proteomic dataset from human synaptosomal P2 fractions of dementia-AD brains, using subcellular fractionation methods similar to ours (59). In the human study, 26 synaptic proteins were significantly increased while 71 decreased in AD synaptosomes, as compared to non-AD controls (59). FET analysis showed that synaptic proteins that decreased in AD, were enriched in mouse M2 synapse/mitochondria (e.g. Rph3a, Ndufs8) module while synaptic proteins that increased in AD (e.g. Dgkg, Uchl1), were enriched in the mouse M1 somatodendritic module (**Fig. 6G**).

Overall, our results integrating mouse Camk2a synaptosome proteomics data with three independent human AD proteomics datasets show that M2 (mitochondria) and M6 (synapse-glutamatergic) modules are (1) preserved across species, (2) strongly linked to AD pathology status and cognitive performance, and (3) highly enriched in pro-resilience proteins. These results nominate molecular targets for future mechanistic validation and therapeutics and link neuroinflammation with mechanisms of cognitive resilience.

## DISCUSSION

The high metabolic demands and rapid molecular events occurring in the neuronal synapse require unique sub-cellular and molecular adaptations of this neuronal compartment. Accordingly, mRNA, mitochondria, lysosomes and translational (ribosomes) machinery are transported to the synapse where they can locally support energy, metabolic and protein synthesis demands (1–3). Neurodegenerative disease processes ultimately cause neurological symptoms by disrupting synaptic function and structure, often starting at early stages that precede overt neurodegeneration or cognitive deficits (7–10, 25). Neuroinflammation, primarily mediated by activated microglia and astrocytes, is an important early and causal mechanism that can lead to synaptic dysfunction and synaptic loss in neurodegenerative disorders including AD (9, 13). Previous proteomic studies using bulk brain homogenates and synapse-enriched fractions (synaptosomes) from post-mortem human samples, longitudinal aging studies and animal models have highlighted the pathological importance of synaptic protein changes in neurodegenerative diseases such as AD, where the loss of synaptic proteins is the strongest correlate with cognitive decline (20, 25, 67–69). Therefore, deciphering the molecular events occurring in neurons and more specifically in their synapses, can provide key mechanistic insights into disease progression and neuronal vulnerability as well as in the nomination of translationally-relevant molecular targets. Since functionally important molecular changes in vulnerable synapses occur at the protein (proteomic) level, it is critical to determine unique proteomic composition of neuronal synapses and examine how synaptic compartment proteomes change in response to disease-relevant processes (e.g. neuroinflammation).

In this study, we have applied cell type-specific *in vivo* biotinylation of proteins (CIBOP) to label Camk2a excitatory neuronal proteomes, and imposed subcellular fractionation to brain lysates, followed by proteomic analysis of biotinylated neuronal proteins. This approach allowed us to obtain the native-state proteome of crude synaptosomes containing presynaptic (mitochondria and synaptic vesicles) and postsynaptic compartments of excitatory neurons, while directly contrasting them with the whole neuronal proteome, from mouse brain. We identified 127 unique synaptic proteins, some expected synaptic proteins (Dlg4, Shank2, Syn3) and ion channels (Grin1, Grin2a, Gria2), as well as several proteins not observed or under-sampled by bulk proteomics approaches. Next, we examined how the synaptic compartment of excitatory neurons responds to neuroinflammatory stress induced by systemic LPS, and contrasted synaptic proteomic alterations with somatodendritic compartmental changes. We observed that the effects of neuroinflammation on the synaptic compartment were indeed distinct from the somatodendritic compartment, suggestive of compartment-specific effects of neuroinflammation. Most of the proteomic effects of LPS on neuronal compartments were also not observed at the bulk tissue proteomic levels, emphasizing the importance of using the CIBOP approach to investigate neuron-specific disease mechanisms *in vivo*. To compliment proteomics data, using an adapted Luminex approach to directly measure neuron-derived phospho-signaling proteins, we found that neuroinflammation leads to suppressed MAPK signaling specifically in the synapse, also not observed in the somatodendritic compartment or the bulk brain lysate. We found that LPS specifically increased mitochondrial protein and pro-resilience protein levels while decreasing synaptic vesicle and cytoskeletal proteins in the synaptosome-enriched neuronal proteome. In support of the functional consequences of these proteomic effects of neuroinflammation, we observed subtle detrimental ultrastructural changes in excitatory asymmetric synapses. Next, we examined how these neuroinflammation-dependent and neuroinflammation-independent proteomic changes in neurons from mouse brain, correspond to observed brain proteomic changes occurring in existing post-mortem human bulk brain and synaptosomal proteomes, as well as with existing markers of cognitive resilience. We found that the mitochondrial and synaptic changes in mouse neurons due to LPS-induced neuroinflammation, were relevant to human AD. Specifically, while neuronal mitochondrial proteins increased with neuroinflammation, these decreased in human AD. Conversely, decreased levels of synaptic structural and vesicle proteins due to neuroinflammation, were also decreased in human AD brain and synaptosomes. Markers of cognitive resilience (e.g. Rph3a) were importantly modulated by neuroinflammation, supporting the relevance of our findings in a mouse model of neuroinflammation with molecular changes occurring in human AD pathology. Taken together, our study extends the CIBOP approach to examine the synaptic compartment *in vivo*, to identify compartment-specific effects of neuroinflammation and lays the foundation for future studies in neurological disease models.

### Unique proteomic characteristics of the excitatory neuronal synaptic compartment

Synaptosomes−isolated nerve terminals containing presynaptic and postsynaptic membranes, mitochondria and synaptic vesicles (70)−are widely used to study synaptic function and molecular characterization of synaptic protein composition or signaling (39). For instance, synaptosome proteomics studies of post-mortem human tissue from AD patients have revealed important insights on synaptic protein composition alterations in disease (59, 67, 71–75). However, post-mortem bulk tissue human studies are certainly limited to disease endpoints and lack cell-type-specific resolution. Since longitudinal ante-mortem molecular studies are not feasible in living individuals, mouse models represent reasonable systems to investigate cell-type-specific synapse diversity (76–78), selective synaptic vulnerability or resilience (7, 25, 38, 39), and early synaptic dysfunction mechanisms (34, 79, 80), with the goal of identifying synaptic targets for disease-modification in neurodegenerative diseases (27).

We employed CIBOP to study excitatory synapses by fusing engineered enzymes (e.g. TurboID) to label cytosolic neuronal proteins, which can then be enriched using affinity capture and characterized by MS (30, 81). Our group recently developed a novel *in vivo* labeling transgenic approach called CIBOP or TurboID-based proximity labeling of proteins coupled with MS for studying native-state proteomes of a specific cell-type (e.g. Camk2a excitatory neurons, PV inhibitory neurons, or Aldh1h astrocytes) (33, 34). Here, we combined Camk2a-CIBOP approach with differential centrifugation to study P2 fractions (crude synaptosomes) and successfully characterized the synaptic compartment proteomes of Camk2a-specific excitatory neurons. Conversely, neuronal proteins differentially higher in the global neuronal proteome, represent the proteome of the somatodendritic compartment of excitatory neurons. Since protein biotinylation from Camk2a-CIBOP mice is neuron-specific, this subcellular fractionation method is advantageous for achieving a higher protein yield compared to density gradient ultracentrifugation and successfully allows the identification and quantification of low-abundance synaptic proteins despite the inherent presence of non-neuronal contaminants (e.g. glial structures) in the crude synaptosomal P2 fractions.

Our Camk2a synaptosome-enriched neuronal proteome (P2-pulldown) showed higher levels of expected synaptic proteins (e.g. Grin1, Grin2a, Grin2b, Dlg4, Shank2, Ncam1, Rab2A, Efhd2, and Kcnma1), while lower levels of cytosolic proteins excluded for synapses and nuclear proteins (e.g. Nefh, Actn1, Tubb2a), consistent with enrichment of synaptic elements including neuronal and synaptic mitochondria seen in our TEM studies. Beyond these expected markers, we identified 127 proteins exclusive to the P2-pulldown proteome that were not seen in the whole neuronal proteome. This synapse-specific group of proteins included mitochondrial (Ndufs2, Vdac2, Sucla2), synaptic (Dlg4, Shank2, Syn3) and ribosomal proteins (Rpl14, Rpl22). The synaptic ribosomal proteins are likely to represent machinery involved in local translation in the synapse (e.g. Rpl18, Rpl29, Rps20). Interestingly, some of these proteins represent key cognitive resilience proteins (e.g. Rph3, Syngap1). The CIBOP approach complements several existing approaches to purify and characterize neuronal synapses from brain tissues.

For instance, Marcassa *et al*. (82), by delivering TurboID constructs via adeno-associated virus injections, characterized the protein composition of the excitatory postsynaptic compartment of two populations of layer 5 neurons in the somatosensory cortex, finding distinct synaptic signatures including proteins regulating synaptic organization and transmission (cell-surface proteins, neurotransmitter receptors and ion channels), and revealing its vulnerability to autism. While we used CIBOP for native-state proteomic profiling of synapses, other isolation-based synaptic approaches have also been developed to assess cell-type-specific synaptic proteome diversity using fluorescent-activated synaptosome sorting (FASS). FASS is a biochemical purification method based on size-gating and fluorescence intensity analysis to sort synaptosomes from transgenic mouse lines that express fluorescent synaptic proteins (37, 38). Pioneer work of Biesemann *et al*. (83), using VGLUT1^Venus^ FASS followed by MS, revealed 163 proteins specifically enriched in isolated glutamatergic synaptosomes, including new synaptic proteins Fxyd6 and Tpd52. Recently, van Oostrum *et al*. (76), using several Cre-inducible knock-in mice, optimized FASS to purify synaptosomes and identified >1,800 unique protein groups characterizing the proteomic signatures of 15 different major synapse subtypes. Both proximity labeling and FASS allow the analysis of cell type-specific synapse protein composition *in vivo.* Though, by using PSD95 to target TurboID, Marcassa *et al*. approach (82) specifically labels and identifies proteins localized to the postsynaptic compartment of excitatory neurons. Instead, CIBOP and FASS successfully enable analysis of all synaptic compartments, but differ in the strategy to identify enriched synaptic proteins. For instance, van Oostrum *et al*. (76) compared fluorescently labeled synaptosome proteomes to an unsorted precursor synaptosome population, determined synapse-enriched proteins by quantitative enrichment, and directly compared proteome diversity across synapse types. In contrast, we compared synaptosome proteomes to bulk brain homogenate proteomes (i.e. somatodendritic compartment) from Camk2a-CIBOP and non-CIBOP mice. Synaptosome preparations are known to be heterogeneous, containing a mixture of different synapse types and non-neuronal contaminants (e.g. glial structures), which might limit the accurate interpretation of proteomics data (83, 84). Notably by using Cre/lox system, our Camk2a-CIBOP approach combined with synaptosome enrichment successfully overcame this major limitation and allowed the identification of excitatory neuron-specific synaptic compartment proteins. Beyond a tool to resolve the proteomes of vulnerable synapses, Camk2a-CIBOP also allowed us to examine how the synaptic compartment of excitatory neurons responds to neuroinflammatory stress induced by systemic LPS, revealing molecular insights into its selective vulnerability, and establishing a foundation for cell-type-specific synapse proteomics work in other preclinical disease models.

### LPS-induced neuroinflammation induces unique excitatory neuron compartment-specific effects

In this study, we used systemic administration of LPS to model neuroinflammation and immune mechanisms, some of which are relevant in AD (15, 61, 85). As expected (63, 86–88), LPS-induced neuroinflammation resulted in body weight loss, microglia activation, and pro-inflammatory cytokine/chemokine release (e.g. Eotaxin and IP-10/Cxcl10). At the bulk proteome level, we found that the effects of LPS on bulk brain homogenates and synaptosomes are molecularly distinct. Particularly in synaptosomes (P2-input), there was an upregulation of actin filament organization, metabolic process and asymmetric synapse-related terms, while a downregulation of electron transfer activity and postsynaptic specialization. The LPS-induced proteomic alterations observed in synaptosomes were in line with previous findings from prefrontal cortex synaptosome samples of LPS-treated mice, reporting LPS-induced proteomic changes in signaling, cytoskeletal, and metabolic processes (89). At the transcriptomic level, similar to previous RNA sequencing studies (16, 63, 88), we found that LPS increased proinflammatory-related gene expression in bulk brain, but transcriptomic changes were more prominent in acutely isolated microglia, consistent with a DAM phenotype signature (16, 90), characterized by downregulation of homeostatic genes, and upregulation of canonical DAM genes (*Clec7a, Cst7, Itgax*), pro-inflammatory genes (*Ccl5, Cxcl10*) and phagocytosis-associated genes (complement *C3*). Complement proteins C3 and C1q are known to tag vulnerable synapses to be engulfed by activated microglia leading to synapse loss in LPS-induced neuroinflammation (63, 91) and transgenic AD mouse models (64, 65). Interestingly, we observed elevated C1q (C1qa, C1qb, and C1qc) protein levels in synaptosomal P2 fractions, indicative of microglia-derived C1q proteins within the synapse. This possibility is supported by recent proteomic findings from crude synaptosomes showing that microglial-derived C1q can be internalized into neurons and accumulates in synaptic terminals, axons and dendrites, where regulates neuronal protein synthesis by interacting with ribonucleoprotein complexes (92). Moreover, we found similar increased C1q protein levels when examined bulk brain proteomes from LPS-treated mice and 5xFAD mice, indicative of common pathological mechanisms between LPS-induced neuroinflammation and AD, including microglial activation and complement C1q production.

At the cell-type-specific proteome level, a key finding of our study is that Camk2a synaptosome (P2-pulldown) proteomes reveal compartment-specific effects of LPS predominantly impacting the synapse, such as an increase in mitochondrial proteins (Pdhb, Ndufc2, Uqcrh) and Rac/RAB GTPases (Rac1, Rab35), while a reduction in cytoskeletal organization (Map4), synaptic (Madd, Snap47) and translation (Eif6) processes. LPS effects on Camk2a synaptic compartment proteomes were profoundly different from those observed in the Camk2a somatodendritic compartment (homogenate-pulldown), highlighting that LPS increased proteins involved in mitochondrial acyltransferases and metabolic activity, while decreased proteins involved in nucleoside triphosphate regulator activity specifically in the synaptic compartment. Another important finding of our study is that LPS reduces cytoskeletal and synaptic proteins. In normal conditions, the cytoskeleton, specifically actin filaments and microtubule-associated proteins (MAPs), plays a crucial role in trafficking and stabilization of mitochondria at the synapse, ensuring ATP supply for synaptic transmission, but also scaffold local translation of synaptic components (93). In our Camk2a synaptosome proteome, we identified Map4 as the most decreased protein in LPS-treated mice. Interestingly, Map4 has been reported to control mitochondrial trafficking by regulating kinesin motor activity *in vitro* (94), suggesting that LPS might disrupt mitochondrial distribution within the synapse.

On the other hand, our Camk2a synaptosome proteomic data extend previous studies (86, 95) reporting that systemic administration of LPS (0.5 mg/Kg, i.p.) induces excitatory synapse alterations. Particularly, when LPS was given for 2 consecutive days, no changes were observed in the number of excitatory synaptic puncta (86), but extended LPS administration for 7 consecutive days significantly decreased vGLUT1 and PSD95 protein levels and excitatory synapse density (95) in the hippocampus. Given that changes in synaptic protein composition can influence synapse morphology and function, we further investigated whether the LPS-induced changes in synaptic proteins lead to observable alterations in the ultrastructure of asymmetric excitatory synapses. In this regard, we found that LPS slightly reduces presynapse size, while increases synaptic vesicle density and PSD length, these subtle morphological abnormalities suggest that LPS might enhance excitatory synaptic transmission. In line with these findings, a recent study using an imaging MS approach correlated with TEM demonstrates that the size of pre- and postsynaptic structures and the number of synaptic vesicles correlate with protein turnover rates (96). Consistently, increased synaptic turnover has been observed in LPS-induced neuroinflammation (91). Additionally, *ex vivo* electrophysiological analysis have shown that LPS can disrupt the balance of excitatory and inhibitory synaptic transmission (63, 86, 95, 97).

Synaptic transmission and plasticity are regulated by signaling mechanisms, including the mammalian target of rapamycin (mTOR) and mitogen-activated protein kinase (MAPK) pathways (4, 5, 98–101). On the one hand, mTOR modulates the activity of translation initiation machinery responsible for synthesizing new proteins at postsynaptic sites, which are essential for long-lasting synaptic plasticity (99–101). On the other hand, MAPK are involved in both gene expression and local synaptic modulation by phosphorylating local substates to control their trafficking and synaptic distribution, including scaffold proteins, cadherin-associated proteins, potassium channels, and metabotropic glutamate receptors, underlying the strength of excitatory synapses (4, 5). Remarkably, using our adapted Luminex assay to directly measure phospho-signaling proteins of interest from Camk2a-CIBOP mice, we found that mTOR signaling activation was not altered, contrasting to MAPK signaling which was increased in the P2 fractions compared to bulk brain homogenates, while LPS preferentially suppressed MAPK signaling (including JNK, MEK1, p38, and ERK1/2) in synaptosome-enriched P2 fractions compared to bulk brain homogenates. While both mTOR (99) and MAPK (4, 5) pathways are recognized as crucial regulators of synaptic plasticity, these findings suggest that LPS-induced neuroinflammation might induce subtle alterations of synaptic function via MAPK-mediated mechanisms. Collectively, our data demonstrate that LPS induces synaptic compartment-specific proteomic changes, alters asymmetric synapse ultrastructure and impairs MAPK signaling in Camk2a synaptosomal fractions, nominating new key proteins and pathways that render selective excitatory synapse vulnerability to neuroinflammatory stress.

### Neuron-specific effects of neuroinflammation on mitochondria

While the crude P2 synaptosome preparation enriches synaptosomes, it also enriches mitochondria, most of which are derived from neurons and astrocytes. The Camk2a-CIBOP approach further enforces neuronal specificity to our analyses, allowing us to investigate how neuron-specific mitochondria are impacted by LPS-induced neuroinflammation. Recent work demonstrated that systemic LPS induces mitochondrial fragmentation in neurons (102) and alters synaptosomal mitochondrial bioenergetics by increasing ATP production (103), but limited information exists regarding LPS effects on synaptic mitochondria proteomes. While the LPS-increased levels of mitochondrial proteins observed in Camk2a synaptosome proteomes might result from alterations in mitochondrial dynamic mechanisms (e.g. enhanced biogenesis, reduced mitophagy, unbalanced mitochondrial fusion-fission, or disrupted anterograde/retrograde transport), it is noteworthy that only few mitochondrial regulators (Dnm1l, Mfn2, and Opa1) were present in our Camk2a synaptosome proteome, without changes in response to neuroinflammatory stress, suggesting that LPS might stimulate synaptic local translation to fulfill the high metabolic demands resulting from alterations in synaptic transmission/plasticity, rather than causing disruptions in synaptic mitochondrial dynamic mechanisms.

Previous studies demonstrated that whole brain synaptic mitochondria have a distinct proteomic profile than non-synaptic mitochondria (104, 105). For example, in comparison to non-synaptic mitochondria, synaptic mitochondria exhibit lower levels of Pdhb (pyruvate dehydrogenase E1 subunit beta) (104), a component of the pyruvate dehydrogenase complex (PDH) essential for energy production in mitochondria. This difference in Pdhb levels might be related to distinct functional demands since synaptic mitochondria are crucial for generating the ATP needed for synaptic transmission and calcium homeostasis, but also might highlight a potential vulnerability of synaptic mitochondria to oxidative damage and calcium overload (105). Interestingly, we observed that LPS increases Pdhb levels and other mitochondrial proteins in Camk2 synaptosomes, which we interpret as a compensatory mechanism for sustaining high metabolic demands consistent with previous findings showing LPS-increased synaptic mitochondrial respiration (103, 106). Similar abnormalities in mitochondrial metabolism were reported to occur early in AD progression. For instance, Dentoni *et al*. (107) showed increased mitochondrial respiration and ATP levels in primary cortical neurons derived from App knock-in mice. Moreover, Naia *et al*. (108) identified an early hypermetabolic phase mainly characterized by increased mitochondrial oxidative phosphorylation (e.g. PDH activity), preceding amyloid-β deposition, autophagy, and synaptic disorganization, in the hippocampus of App knock-in mice. Recently, using PV-CIBOP approach, our group characterized the PV interneuron proteomes in 3 month-old 5xFAD mice (minimal plaque deposition), revealing unique signatures of increased mitochondria and metabolism as one of the most significantly altered pathways at early stages of amyloid-β pathology, changes not resolved in the bulk brain proteome, enlightening selective vulnerabilities of PV interneurons to neurodegeneration in AD (34). Strikingly, Pdhb, one of most increased proteins in Camk2a-synaptosomes from LPS-treated mice, was reported to be increased at early-stage AD, but decreased at advanced-stage AD (109). These observations suggest that LPS-induced effects on mitochondria overlap with neuronal events in early-stage AD triggered by activated microglia, but disease progression eventually overcomes microglial protective mechanisms, leading to chronic neuroinflammation, oxidative stress, excessive pro-inflammatory cytokine production, synapse engulfment, blood-brain barrier compromise, and plaque deposition.

Further studies are needed to determine whether these early mitochondrial changes play a protective or detrimental role in subsequent bioenergetic failure and mitochondrial dysfunction, including glucose hypometabolism and alterations in several mitochondrial enzyme activities, oxidative stress, calcium ion imbalance, and mitochondria dynamics alterations observed in late-stage AD (110, 111). However, alterations in mitochondrial protein levels might go beyond impacting neuronal energy production, but also significantly contributing to synaptic dysfunction and signaling impairments, ultimately causing synaptic loss and cognitive decline in AD. Understanding the complex interplay between early mitochondrial protein changes and synaptic dysfunction is crucial for developing effective AD therapies. While emerging mitochondria-targeted therapies such as mitochondrial gene therapy and pharmacological interventions (e.g. antioxidants, mitochondrial biogenesis enhancers) focused on restoring mitochondrial function show promise to slow down AD progression in preclinical studies (112–114), more research is needed to translate these therapies into effective treatments for humans given the current challenges concerning off-target effects and delivery mechanisms. For example, chronic treatment with a mitochondrial complex I inhibitor starting at presymptomatic stage (2.5 month-old) in APP/PS1 mice improved cognitive functions and synaptic strength, enhance metabolic function (increased glucose oxidation and ATP preservation), and mitigated oxidative stress and inflammation, through selective activation of mitochondria-associated AMPK pathway, suggesting its potential as a neuroprotective agent by targeting mitochondrial dysfunction at early-stage AD (115). By using a neuron-specific proteomic approach combined with synaptosome enrichment, our findings identify specific neuronal mitochondrial proteins impacted by LPS within the synaptic compartment, enabling future studies focused on the development of mitochondrial-targeted therapies in preclinical neurodegenerative disease models.

### Excitatory neuron synaptic molecular signatures are associated with cognitive resilience in humans

Looking for relevance of Camk2a synaptosome molecular signatures in human AD, we assessed existing human post mortem bulk brain proteomic data from controls and AD cases to identify AD-related protein co-expression modules (21). Interestingly, this integrative analysis identified shared and divergent molecular changes between mouse LPS-neuroinflammation model and human AD. Specifically, LPS increased modules M2 and M5 enriched in mitochondrial proteins, while decreased module M9 Magenta enriched in pre-synaptic vesicle and GABAergic transmission. In contrast to Camk2a-synaptosomes, mitochondria-enriched module was less abundant in AD cases compared to both asymptomatic AD and Control groups. Given mitochondrial alterations in AD are age- and disease-stage-dependent (20, 105, 108, 109, 116, 117), this discrepancy suggests that the LPS-neuroinflammation model recapitulates mitochondrial alterations observed at early-stage AD mouse models (34, 108), potentially driven by amyloid-β. On the other hand, synaptic-enriched modules decreased in LPS-neuroinflammation model and human AD. Among commonalities, the synaptosome associated protein 47 (Snap47), located at the postsynaptic compartment and involved in AMPA receptor trafficking (118), is one of the LPS-decreased proteins in Camk2a synaptosome proteomes. Consistently in AD, Snap47 levels decreased, among other synaptic proteins, correlating with the severity of cognitive decline (24, 67, 116). Besides AD pathology status, eigen proteins from synaptic-enriched modules impacted by LPS treatment were strongly linked to cognitive performance in AD (21). Taking into consideration the complexity of AD pathogenesis, LPS-induced neuroinflammation effectively model cellular and molecular mechanisms of mitochondrial and synaptic alterations seen in early-stage AD, but does not completely recapitulate the phenotype for mitochondrial dysfunction observed in late-stage AD.

As additional measures for assessing the relevance of mouse data to human, we integrated our mouse module proteomics data with molecular signatures of cognitive resilience to AD. Yu *et al*. (22) define proteins for cognitive resilience as targets that may be associated with slowing or preventing cognitive decline regardless of the presence, number, or combination of common neuropathologic conditions. In the human synaptosome study, Carlyle *et al*. (59) analyzed P2 fractions from 100 AD subjects and controls. Similar to the cognitive resilience study (22), but opposed to our Camk2a synaptosome findings from LPS-treated mice, the mitochondria-enriched M2 Blue module was decreased in the human synaptosomal fractions from dementia-AD individuals (59). Interestingly, the M2 module contains multiple pro-resilience proteins, such as Ndufs8, Atad1 and Rph3a. Ndufs8 is a subunit of electron transport chain complex I involved in mitochondrial respiration (119), the ATPase Atad1 plays a role in maintaining mitochondrial function by facilitating the degradation of mis-localized or misfolded proteins (120), while Rph3a is crucial for retaining NMDA receptors at excitatory synaptic sites (121). Among the resilience proteins from our postsynapse-enriched M6 module, there is Rab35, which regulates protein trafficking and amyloid-β production (122), and Gria3, a protein subunit for AMPA receptors, known to be decreased in advanced AD (109). Unexpectedly, the M1 module, containing anti-resilience proteins related to bioenergetic and synaptic dysfunction (Dgkg, and Tkt, Uchl1), was increased in synaptosomes from AD individuals, while no changes were identifiable in response to LPS. These findings on Camk2a synaptosomes from LPS-treated mice are in line with our previous study in 5xFAD mice (34), where PV-interneuron proteomic signatures revealed increased mitochondria and metabolism related proteins, but reduced synaptic and cytoskeletal proteins at early-stage of AD pathology, and remarkably, these PV-specific changes were also not captured by bulk brain proteomes. Together these data suggest that early in the disease, vulnerable excitatory and inhibitory neurons might initially try to compensate for the detrimental impact of amyloid-β pathology on their synapses through mechanisms such as upregulation of mitochondrial proteins. However, as the disease progresses, vulnerable neurons might experience a decline in synaptic pro-resilience proteins, leading to synaptic dysfunction and ultimately contributing to synapse loss and neurodegeneration.

## Limitations of this study

First, we acknowledge that crude synaptosomes (P2 fraction) might contain non-synaptic neuronal and glial structures, mostly membrane fragments and extra-synaptosomal mitochondria (84). However, an advantage of using crude synaptosomes from neuron-CIBOP mice, rather than preparing a “purer” synaptic fraction via density gradient centrifugation (41), is the ability of identifying biotinylated proteins specifically within neurons without the need of further glial contaminant depletion, while maintaining a high protein yield for downstream analysis. Second, to ensure that observed changes in specific mitochondrial proteins were due to LPS and not to changes in mitochondrial mass (i.e. total mitochondria protein content), we used mitochondrial abundance-based normalization. After applying this normalization, we confirmed that the key mitochondrial DEPs (e.g. Pdhb, Ndufc2, Uqcrh) increased by LPS remained unchanged, suggesting that the observed changes were not due to a global increase in mitochondrial mass. Therefore, we concluded that LPS induces specific changes in the intrinsic properties of the neuronal mitochondria, rather than affecting their overall quantity at the excitatory synapses. Third, even though Camk2a-Cre-ert2 mice have been widely used to target excitatory glutamatergic neurons, it is important to acknowledge that Camk2a expression can occur at low levels in non-excitatory neurons (123). Lastly, we acknowledge that our analysis of P2 fractions from whole brains might have masked potential LPS effects arising from selective vulnerable brain regions to neuroinflammation. Given the vast molecular diversity of neurons (124) and their synapses (76–78), and rapid technical advances in MS paradigms (125), future studies should also transition from data-dependent acquisition (DDA) to data-independent acquisition (DIA) methods, which offers a higher sensitivity and a broader protein coverage (37, 126), potentially leading to quantification of low-abundance proteins within neurons and their synapses towards a more comprehensive profiling of subtle proteomic changes induced by neuroinflammation.

## Conclusion

In conclusion, our neuron-CIBOP combined with differential centrifugation is a powerful experimental approach to investigate synaptic compartment-specific molecular changes occurring at the proteomic level under native conditions. The Camk2a synaptosome proteomic signature was enriched in mitochondrial, synaptic transmission, and ribosomal proteins. Our study reveals unique effects of LPS-neuroinflammation on excitatory synapse proteomic composition, ultrastructure, and signaling, while highlighting important commonalities and differences between the molecular mechanisms involved in LPS-neuroinflammation and AD pathogenesis. Looking forward, the complexity of mitochondrial and synaptic dysfunction mechanisms in neurodegenerative diseases, proteomic profiling of cell-type-specific synaptosomes via CIBOP combined with differential centrifugation might offer valuable molecular insights into the protein composition changes occurring at different ages in vulnerable synapses in AD mouse models for better understanding disease progression and developing effective disease-modifying therapies.

## Supporting information

Analyses and datasets related to Fig. 1

Analyses and datasets related to Fig. 2

Analyses and datasets related to Fig. 3

Analyses and datasets related to Fig. 4

Analyses and datasets related to Fig. 5

Analyses and datasets related to Fig. 6

Analyses and datasets related to Supplementary Fig. S1

Analyses and datasets related to Supplementary Fig. S2

Analyses and datasets related to Supplementary Fig. S3

Analyses and datasets related to Supplementary Fig. S4

Analyses and datasets related to Supplementary Fig. S5

## Abbreviations

AD: Alzheimer’s disease
BSA: Bovine serum albumin
Camk2a: Calcium/calmodulin dependent protein kinase II Alpha
CIBOP: Cell type-specific In vivo Biotinylation of Proteins
DAM: Disease-associated microglia
DEPs: Differentially expressed proteins
DTT: Dithiothreitol
D-PLSR: Discriminant partial least squares regression analysis
FDR: False discovery rate
GFAP: Glial fibrillary acidic protein
GSVA: Gene set variation analysis
HSP60: Heat shock protein 60
Iba1: Ionized calcium-binding adaptor molecule
1 IP-10: Interferon gamma-induced protein 10
IPA: Ingenuity Pathway Analysis
LFQ-MS: Label-free quantitative mass spectrometry
LPS: Lipopolysaccharide
LV1: Latent variable one
MAPK: Mitogen-activated protein kinases
MAPs: Microtubule-associated proteins
mTOR: Mammalian target of rapamycin
NeuN: Neuron specific nuclear protein
NfL: Neurofilament light chain
PCA: Principal-component analysis
PDH: Pyruvate dehydrogenase complex
Pdhb: Pyruvate dehydrogenase E1 subunit beta
PFA: Paraformaldehyde
PSD: Postsynaptic density
PSD95: Postsynaptic density protein 95
PWAS: Proteome-Wide Association Study
SDHA: Succinate dehydrogenase complex flavoprotein subunit A
SVs: Synaptic vesicles
SV2A: Synaptic vesicle glycoprotein 2A
SynGO: Synaptic gene ontology
TEM: Transmission electron microscopy
TFA: Trifluoroacetic acid
TMT: Tandem Mass Tag
VDAC: Voltage-dependent anion channel
VGLUT1: Vesicular glutamate transporter 1
WGCNA: Weighted gene co-expression network analysis
WT: Wild type

## DECLARATIONS

### Ethics approval and consent to participate

Not applicable.

### Consent for publication

Not applicable.

### Availability of data and materials

The MS proteomics data have been deposited to the ProteomeXchange Consortium and can be accessed via the PRIDE data repository with the dataset identifier PXD068874.

## Authors’ contributions

SR and CE-G conceptualized and designed the study. CE-G, US, PK, DK, SM, BRT, SS, HX, LC, PB, DMD, NTS, LBW, VF, and SR conducted experiments, analyzed data and interpreted the results. US and SR analyzed mass spectrometry data. TJW and XL processed samples for transmission electron microscopy analysis and acquired electron micrographs. CE-G, US and SR wrote the original draft of the manuscript. All authors reviewed, provided critical feedback and approved the final version of the manuscript. NTS, LBW, VF and SR acquired funding for the studies. LBW, VF and SR provided laboratory resources for the experiments.

## Acknowledgements

Research reported in this publication was supported in part by Emory University subsidized cores Emory Integrated Proteomics Core (EIPC, RRID:SCR_023530) and Integrated Electron Microscopy Core (IEMC, RRID:SCR_023537). Additional support was provided by the Georgia Clinical & Translational Science Alliance of the National Institutes of Health under Award Numbers UL1TR002378 (EIPC) and UL1TR000454 (IEMC), and the National Institutes of Health under award numbers P30CA138292 and S10RR025679. The content is solely the responsibility of the authors and does not necessarily represent the official views of the National Institutes of Health.

