## Supplementary Figures and legends for "Neuroinflammatory Stress Preferentially Impacts Synaptic MAPK Signaling and Mitochondria in Excitatory Neurons"

**ADDITIONAL FILE 12. Supplementary figures and legends**

Supplementary Figures: 5

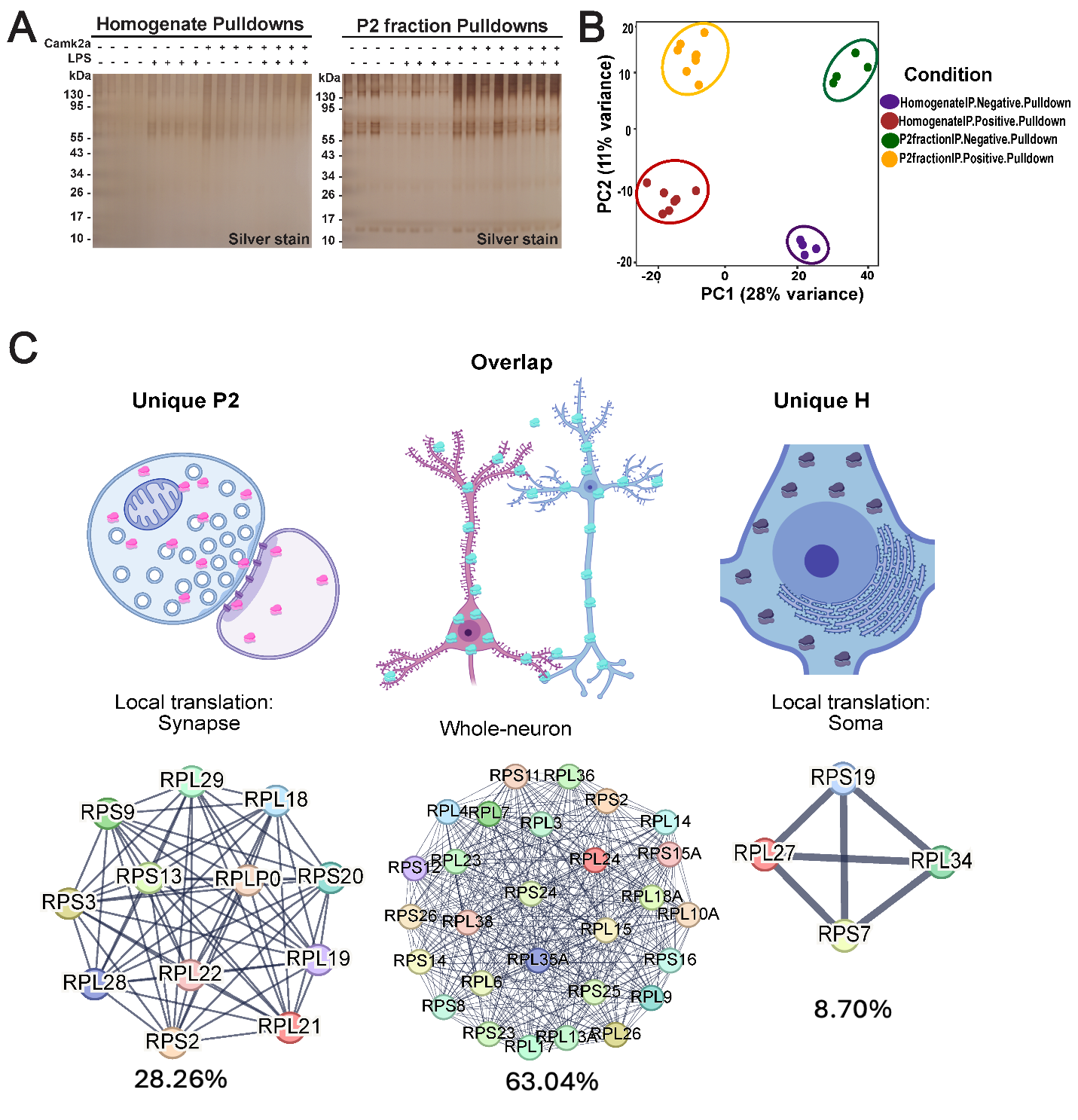

**Supplementary Figure S1. Validation of excitatory synapse-specific proteomics using Camk2a-CIBOP and P2 fractionation. (A)** Silver stain of streptavidin affinity Homogenate and P2 fraction pulldown samples confirm strong protein biotinylation in Camk2a-CIBOP mice as compared to limited biotinylation in non-CIBOP negative controls. **(B)** PCA of MS data from streptavidin affinity enriched proteomes showing Camk2a homogenate-pulldown and Camk2a P2-pulldown proteomes clustered away from non-CIBOP negative controls. **(C)** Localization of ribosomal proteins to subcellular compartments. Our analysis revealed that 28.26% (12) of the proteins were uniquely present in the P2 fraction, 8.70% (4) were uniquely localized in the homogenate fraction, and 63.04% (28) were shared between the homogenate and P2 fractions. First, the ribosomal proteins uniquely identified in the P2 fraction, were enriched in pathways related to ribosome assembly, initiation of translation, and communication between the 40S and 60S ribosomal subunits. Second, the proteins uniquely localized to the homogenate are associated with the 60S ribosomal subunit and are critical for peptide bond formation and polypeptide chain elongation. Third, the overlapping set included essential proteins integral to both the small and large ribosomal subunits, playing central roles in ribosome function and cellular translation. Also see Additional file 7 for related analyses and datasets. Image was created using BioRender.com.

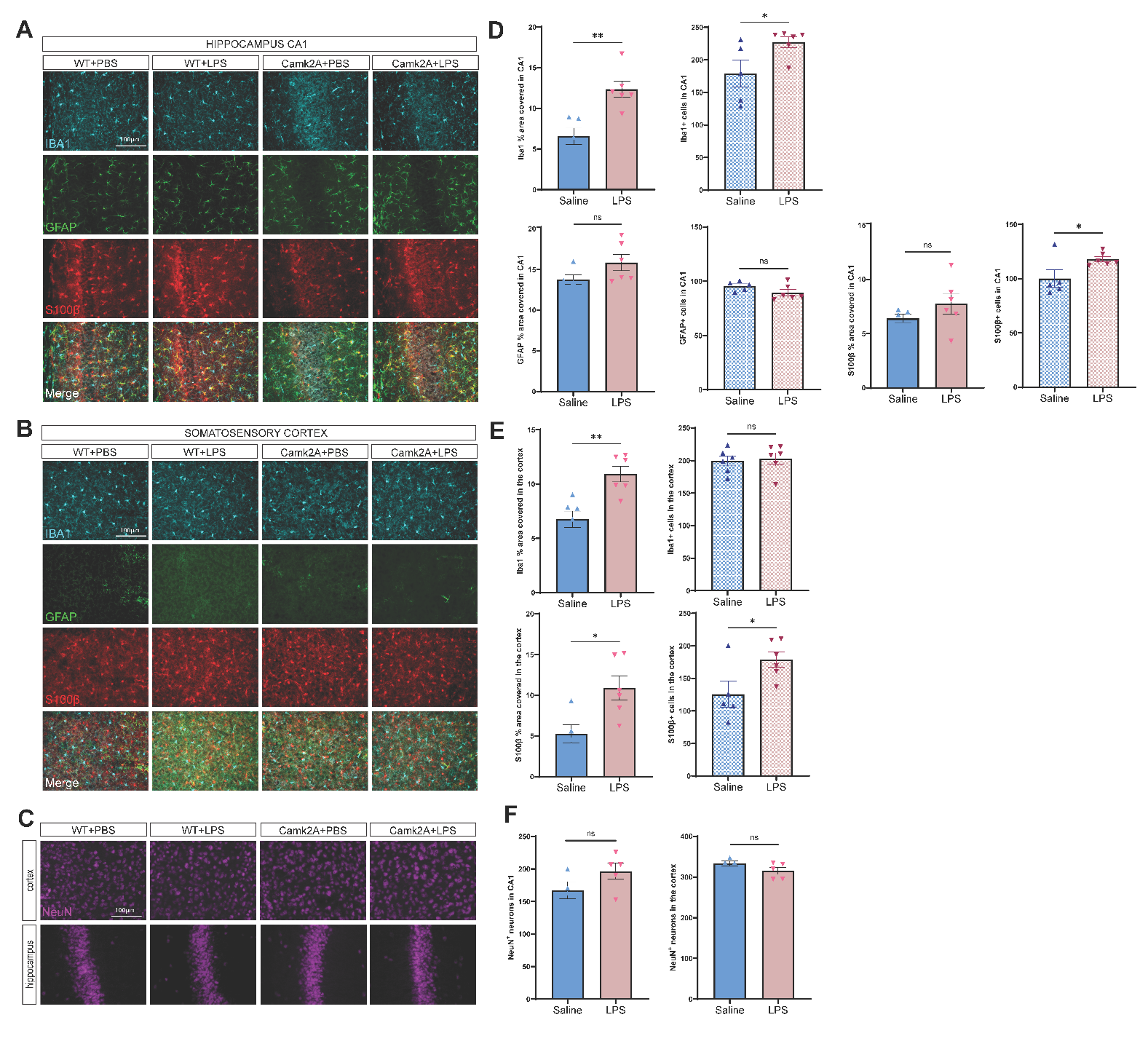

**Supplementary Figure S2.** **Systemic LPS administration induced microglial activation and astrocyte reactivity.** **(A and D)** In hippocampal CA1, LPS-treated mice displayed a higher percentage area covered by Iba1, as well as increased number of Iba1^+^ and S100β^+^ cells, indicative of microglia and astrocyte activation. **(B and E)** Similarly, in the somatosensory cortex, LPS-treated mice displayed a higher percentage area covered by Iba1 and S100β, while GFAP^+^ astrocytes were barely detected in this brain area. **(C and F)** No evidence of neuronal loss was observed in LPS-treated mice. For each marker, a mean ± standard error of the mean (SEM) was calculated for each group (*n* = 4–6). Data were analyzed by unpaired t-test for comparison between Saline- and LPS-cohorts. Also see Additional file 8 for related analyses and datasets.

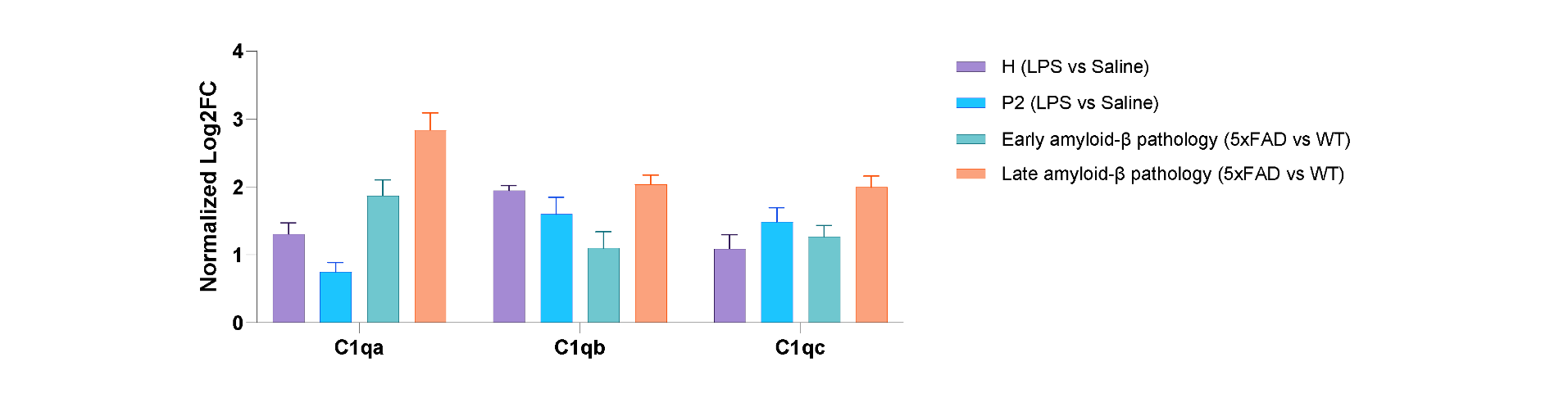

**Supplementary Figure S3. Complement system protein induction in** **LPS-treated WT mice compared to 5xFAD mice.** For LPS-treated WT mice Log2FC C1q values were normalized to saline-treated WT mice, while for 5xFAD mice, Log2FC C1q values were normalized to age-matched WT mice. Elevated levels of C1qa, C1qb, and C1qc in homogenate-inputs and P2 fraction-inputs of LPS-treated WT mice were similar to levels measured in bulk brain homogenates of 5xFAD mice with early or late amyloid-β pathology (*n =* 4‒6/group). Also see Additional file 9 for related analyses and datasets.

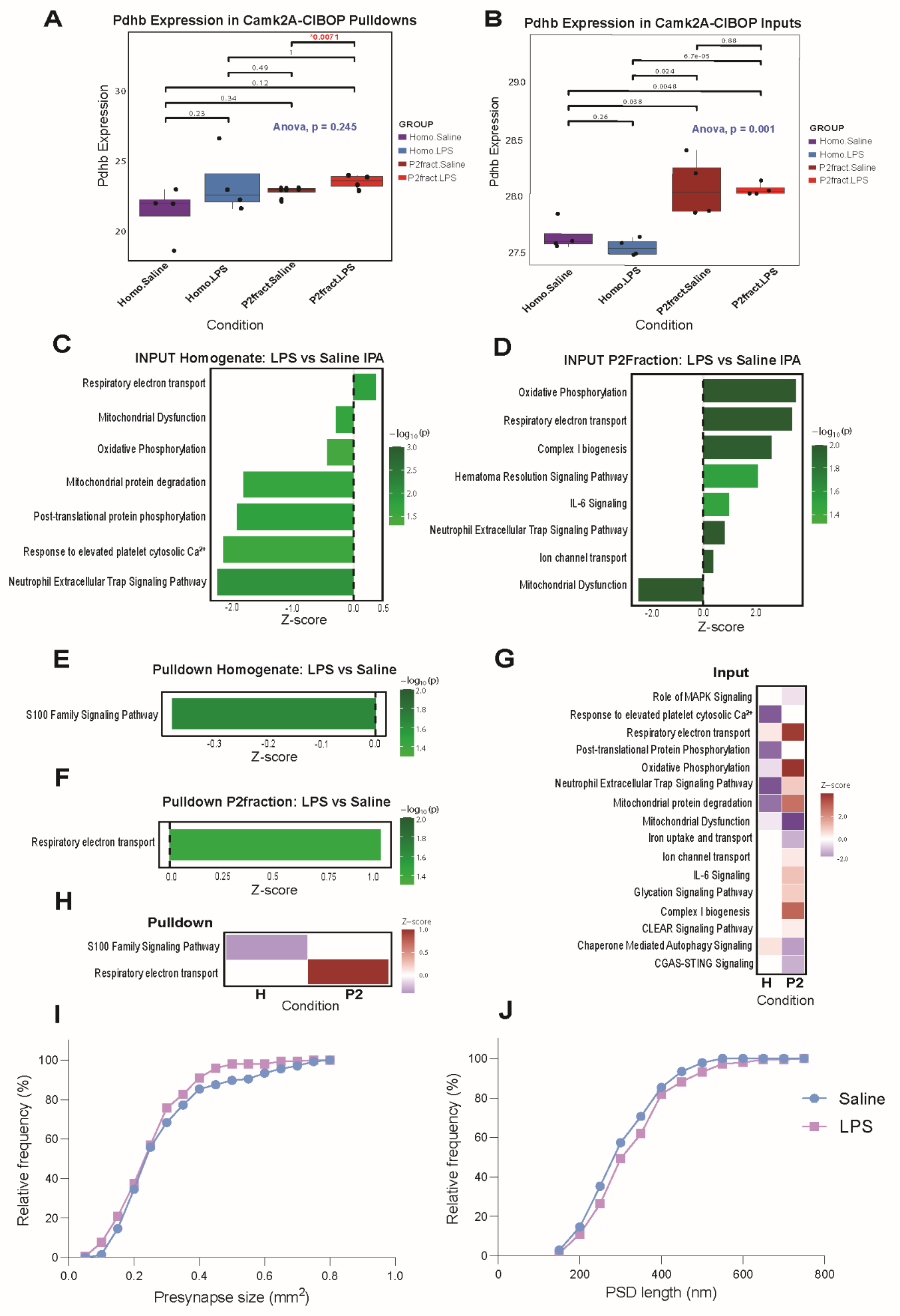

**Supplementary Figure S4. LPS increased Pdhb expression in Camk2a-CIBOP P2 synaptosomal fraction.** Box plots showing Pdhb expression in Camk2a-CIBOP **(A)** pulldown samples and **(B)** input samples. Expression values are quantile normalized and presented as normalized expression levels. Box plots display median (line), interquartile range (box), and full range excluding outliers (whiskers). Individual data points are shown as dots. One-way ANOVA, with individual pairwise comparisons shown; p-values. Notably, Pdhb expression was uniquely elevated in Camk2a-CIBOP P2 synaptosomal fraction compared to homogenate and saline controls. **(C)** Significant canonical IPA pathways induced by LPS in homogenate-input, **(D)** P2-input, **(E)** homogenate-pulldown **(F)** P2-pulldown. **(G)** Comparative heatmap for IPA pathways induced by LPS in homogenate-input compared to P2-input. **(H)** Comparative heatmap for IPA pathways induced by LPS in homogenate-pulldown compared to P2-pulldown. Cumulative distribution curves for **(I)** presynaptic bouton size and **(J)** PSD length of asymmetric synapses. Saline *n* = 136 and LPS *n* = 144 asymmetric synapses from three WT mice for each treatment. There was no significant difference between the two cumulative distribution curves [K-S test, presynapse size: D=0.09273, P=0.5845; PSD length: D=0.1401, P=0.1283]. Also see Additional file 10 for related analyses and datasets.

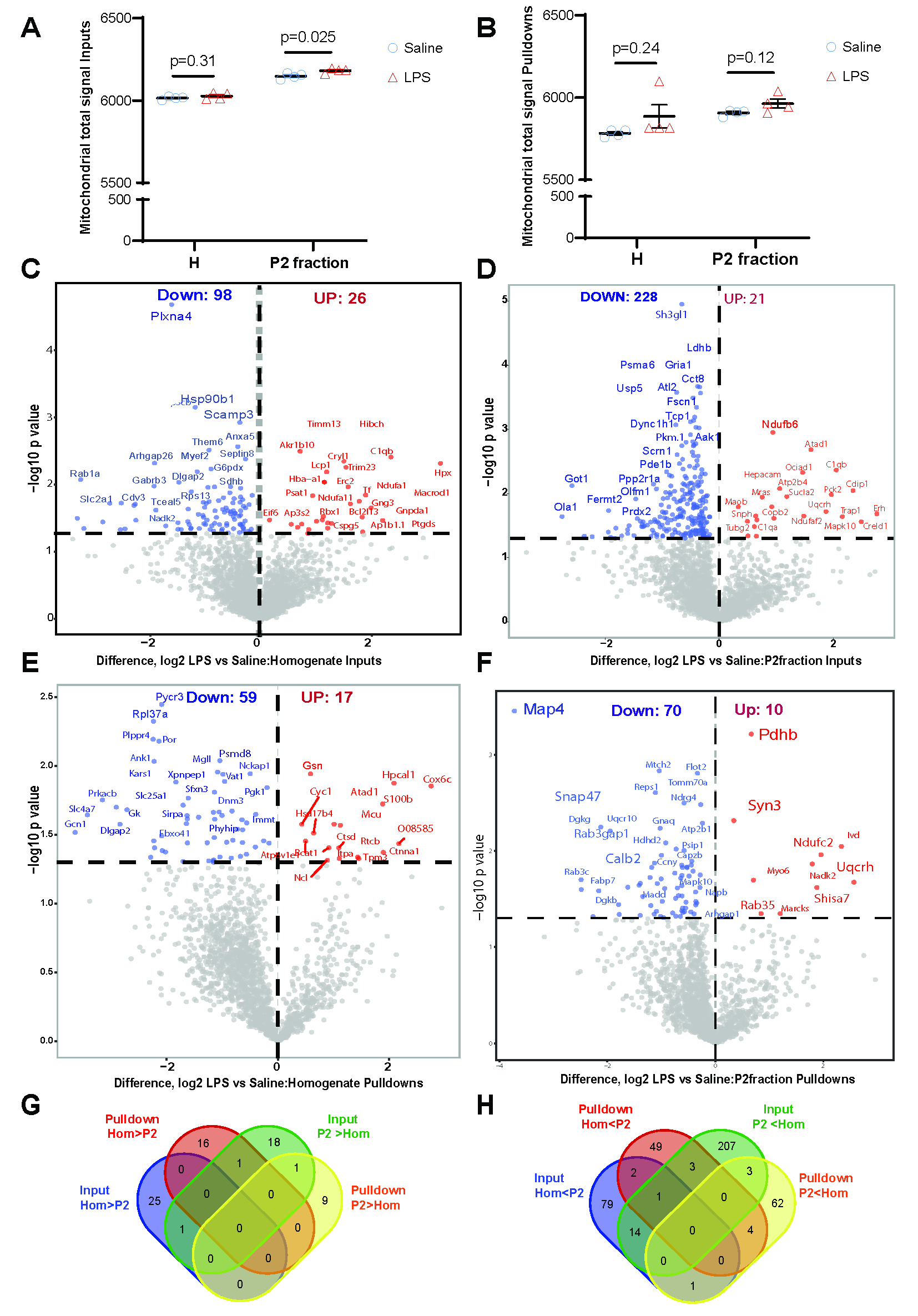

**Supplementary Figure S5. Validation of LPS-induced DEPs after mitochondrial abundance-based normalization.** Mitochondria total signal in **(A)** Inputs and **(B)** Pulldowns. Data were analyzed by unpaired t-test for comparison between Saline- and LPS-cohorts. Volcano plots showing the effects of LPS on **(C)** bulk brain proteomes (LPS homogenate-input vs Saline homogenate-input) and on **(D)** P2 synaptosomal fraction proteomes (LPS P2-input vs Saline P2-input). Volcano plots showing the effects of LPS on **(E)** Camk2a whole-neuron proteome (LPS homogenate-pulldown vs Saline homogenate-pulldown) and **(F)** Camk2a synaptosome proteome (LPS P2-pulldown vs Saline P2-pulldown). Venn diagrams of the unique and shared proteins that were identified from homogenate and P2 fraction inputs and pulldowns **(G-H)**. Additional file 11 for related analyses and datasets.
